# Temporal dynamics of neural responses in human visual cortex

**DOI:** 10.1101/2021.08.08.455547

**Authors:** Iris I.A. Groen, Giovanni Piantoni, Stephanie Montenegro, Adeen Flinker, Sasha Devore, Orrin Devinsky, Werner Doyle, Patricia Dugan, Daniel Friedman, Nick Ramsey, Natalia Petridou, Jonathan Winawer

**Affiliations:** Department of Psychology, New York University, New York, NY 10003, USA; Video & Image Sense Lab, Institute for Informatics, University of Amsterdam, 1098 XH Amsterdam, The Netherlands; Department of Neurology and Neurosurgery, Utrecht Brain Center, University Medical Center Utrecht, 3584 CX, Utrecht, The Netherlands; Department of Neurology, New York University Grossman School of Medicine, New York 10016, NY, USA; Department of Neurosurgery, New York University Grossman School of Medicine, New York 10016, NY, USA; Department of Psychiatry, New York University Grossman School of Medicine, New York 10016, NY, USA; Department of Radiology, Center for Image Sciences, University Medical Center Utrecht, 3584 CX, Utrecht, The Netherlands

## Abstract

Neural responses to visual stimuli exhibit complex temporal dynamics, including sub-additive temporal summation, response reduction with repeated or sustained stimuli (adaptation), and slower dynamics at low contrast. These phenomena are often studied independently. Here, we demonstrate these phenomena within the same experiment and model the underlying neural computations with a single computational model. We extracted time-varying responses from electrocorticographic (ECoG) recordings from patients presented with stimuli that varied in contrast, duration, and inter-stimulus interval (ISI). Aggregating data across patients yielded 98 electrodes with robust visual responses, covering both earlier (V1-V3) and higher-order (V3a/b, LO, TO, IPS) retinotopic maps. In all regions, the temporal dynamics of neural responses exhibit several non-linear features: peak response amplitude saturates with high contrast and longer stimulus durations; the response to a second stimulus is suppressed for short ISIs and recovers for longer ISIs; response latency decreases with increasing contrast. These features are accurately captured by a computational model comprised of a small set of canonical neuronal operations: linear filtering, rectification, exponentiation, and a delayed divisive normalization. We find that an increased normalization term captures both contrast- and adaptation-related response reductions, suggesting potentially shared underlying mechanisms. We additionally demonstrate both changes and invariance in temporal response dynamics between earlier and higher-order visual areas. Together, our results reveal the presence of a wide range of temporal and contrast-dependent neuronal dynamics in the human visual cortex, and demonstrate that a simple model captures these dynamics at millisecond resolution.

**Significance Statement:** Sensory inputs and neural responses change continuously over time. It is especially challenging to understand a system that has both dynamic inputs and outputs. Here we use a computational modeling approach that specifies computations to convert a time-varying input stimulus to a neural response time course, and use this to predict neural activity measured in the human visual cortex. We show that this computational model predicts a wide variety of complex neural response shapes that we induced experimentally by manipulating the duration, repetition and contrast of visual stimuli. By comparing data and model predictions, we uncover systematic properties of temporal dynamics of neural signals, allowing us to better understand how the brain processes dynamic sensory information.

## Introduction

The manner in which neural responses change over short time-scales, from milliseconds to seconds, is important for understanding cognitive, perceptual, and motor functions. Neural dynamics are critical for decision-making (Gold and Shadlen, 2007; Wang, 2012), motor planning (Churchland et al., 2012), and perception (Heeger, 2017), and are important for achieving improved performance in artificial neural network models (Kubilius et al., 2019; Spoerer et al., 2020). Even for simple static stimuli, neural responses in sensory cortex exhibit interesting and complex temporal dynamics. For example, neural responses in visual cortex start to decrease when a static visual stimulus is prolonged in time (subadditive temporal summation), reduce to stimuli that are repeated (adaptation), and rise less rapidly for low contrast stimuli (phase delay).

Studies of temporal dynamics in visual cortex typically measure neural responses with a tailored set of stimuli designed to investigate one particular kind of temporal dynamics. For example, one study might show visual stimuli that vary in duration to investigate how neural activity sums over time (e.g., Tolhurst et al., 1980), while another study might show repeated stimuli to investigate neural response reductions due to adaptation (e.g., Motter, 2006), and yet another study might vary the level of stimulus contrast to study how input strength affects the rise and fall of neural responses over time (e.g., Albrecht et al., 2002). These phenomena are also often studied with different measurement techniques and different computational models.

Even when the same computational model is used across different stimulus manipulations, model parameters may be fit separately to different experiments (e.g., Zhou et al., 2019), leaving open the question of whether a single model, with a single set of parameters, can *simultaneously* account for the multiple phenomena. Here we investigated multiple temporal dynamics in a single dataset by measuring neural responses to visual stimuli that varied systematically in three different ways: duration, repetition, and contrast. We measured electrocorticographic (ECoG) recordings of human visual cortex, which track neural responses at the millisecond scale with high spatial and temporal precision. From each electrode and for each stimulus, we extracted the time-varying broadband power (50-200 Hz). Using different stimulus manipulations in the same electrodes allows us to investigate links between phenomena and ask whether they can be explained by the same computational mechanism. For example, we demonstrate that reducing stimulus contrast and repeating a stimulus result in surprisingly similar changes in temporal dynamics of neural population responses, and link these changes to specific computational model components.

By mapping the electrodes to a probabilistic retinotopic atlas within each participant and then aggregating measurements over multiple participants, we collected a large, comprehensive sample of neural responses from multiple visual areas in human visual cortex, covering earlier (V1-V3a/b) and higher-order (LO, TO, IPS) retinotopic maps. Testing the same stimuli in the same participants across multiple visual areas clarifies the qualitative similarities across areas as well as the quantitative differences. For example, our approach allows us to address discrepant claims about temporal window length in the visual hierarchy, as there is some evidence both in support of (Hasson et al., 2008; Weiner et al., 2010; Honey et al., 2012), and against (Fritsche et al., 2020) the claim that temporal window length increases along the visual hierarchy.

The paper is structured as follows. We first examine the different temporal dynamics resulting from each stimulus manipulation (changes in duration, repetition and contrast) and their corresponding non-linear effects in area V1. We show that these modulations are well captured by a dynamic normalization model fit simultaneously to all stimulus types, and perform a systematic comparison with reduced versions of the model, demonstrating the contribution of each individual canonical computation to its ability to fit the data. Finally, we investigate to what extent these temporal dynamics vary both across and within visual areas.

## Materials and Methods

### Subjects

Data were measured from 11 participants (6 females) who were undergoing subdural electrodes implantation for clinical purposes. 8 participants (5 females) were included in the dataset after data preprocessing (see below for details). Data from 9 participants were collected at New York University Grossman School of Medicine, New York, USA, and 2 participants were tested at the University Medical Center Utrecht (UMCU), Utrecht, The Netherlands. Written informed consent to participate in this study was given by all the patients. The study was approved by the NYU Grossman School of Medicine Institutional Review Board and the ethical committee of the UMCU, in accordance with the Declaration of Helsinki (2013). All participants were implanted with standard clinical subdural grid and/or strip electrodes. Several NYU participants were additionally implanted with standard clinical depth electrodes. For most participants, electrode implantation and location were guided solely by clinical requirements. Two NYU participants consented to the additional placement of a small, high-density grid (PMT Corporation), which provided denser sampling of underlying cortex. Detailed information about each participant and their implantation is provided in **Table 1** and in **Extended Data Figure 1-1 and 1-2**.

**Table 1:** Overview of patient data included in this dataset. Columns refer to the following: *Subject*: Subject code in dataset. *Age*: Age of patient at time of recording in years. *Sex*: Patient sex (M = Male, F = Female). *Site*: hospital in which recording took place (NYU: NYU Langone Hospital; UMCU: University Medical Center Utrecht). *Trials*: Number of trials collected per condition. *Hemi*: implanted hemisphere (L = left, R = right). *Implantation*: type of electrodes implanted (grid = standard clinical grid; strip = standard clinical strip; depth = depth electrodes; HD grid = high-density grid). *Coverage*: approximate overview of visual cortex covered (lat = lateral, med = medial, vent = ventral, par = parietal, occ = occipital, temp = temporal, ant = anterior, post = posterior; see **Extended Data Figure 1-1** for visualizations per patient). *#Visual*: number of electrodes that had a match with either one of the retinotopic atlases used for initial electrode selection (see Methods and Materials). *Matching areas*: matched retinotopic maps according to the maximum probability atlas by Wang et al., (2014). This includes depth electrodes which were not analyzed in the current study. *#Included*: number of electrodes included in the final dataset, after rejection of depth electrodes, epoch and electrode selection (see Methods and Materials). Patients for whom none of the electrodes survived selection are indicated in gray. Note that while these patients do not contribute to the results reported in this paper, we include these patients in the overview because alternative selection data pruning and electrode selection methods on the publicly available data may yield other inclusion results.

| Subject | Age | Sex | Site | Trials | Hemi | Implantation | Coverage | # Visual | Matching areas | # Included |
| --- | --- | --- | --- | --- | --- | --- | --- | --- | --- | --- |
| P01 | 30 | F | UMCU | 12 | L | strip, depth | lat-occ, lat-par, vent-temp | 9 | TO1, LO1, PHC2 | 0 |
| P02 | 18 | F | UMCU | 6 | R | grid, strip | med-occ, lat-occ, lat-temp, vent-temp | 24 | V1v, V1d, V2v, V3v, V3a, V3d, IPS0 | 9 |
| P03 | 27 | M | NYU | 12 | L | grid, strip, depth | lat-par, ant-temp, post-vent, frontal | 24 | V2d, V3d, V3a, V3b, TO1, IPS0-3, SPL1 | 4 |
| P04 | 28 | M | NYU | 10 | R | grid, strip, depth | lat-occ, ant-temp, frontal | 6 | V3d, V3a, TO1 | 2 |
| P05 | 31 | F | NYU | 12 | R | 2x grid, strip, depth | lat-occ, lat-par, frontal, ant-temp | 40 | V2v, V3v, V3d, V3b, LO1, LO2, TO1, IPS2-3, hV4, PHC1 | 3 |
| P06 | 19 | F | NYU | 12 | L | grid, strip, depth | med-occ, lat-occ, lat-par, ant-temp, frontal | 26 | V1d, V2v, V3d, V3b, hV4, TO1, IPS0, PHC2 | 2 |
| P07 | 41 | F | NYU | 24 | L | grid, strip, depth | lat-occ, lat-par, med-occ, ant-temp, frontal | 17 | V1v, V1d, V2d, V3d, V3a, IPS0-3, FEF | 7 |
| P08 | 47 | M | NYU | 12 | R | grid, strip, depth | lat-par, ant-temp | 4 | LO2, TO1 | 0 |
| P09 | 25 | M | NYU | 24 | L,R | bilateral grid, strip, depth | lat-occ, lat-par | 8 | TO1, PHC2 | 0 |
| P10 | 23 | M | NYU | 36 | R | grid, HD grid, strip, depth | lat-occ, lat-par, post-vent | 123 | V2d, V3d, V3a, V3b, LO1, LO2, TO1, IPS0-2 | 45 |
| P11 | 19 | F | NYU | 36 | L | grid, HD grid, strip, depth | lat-occ, lat-par, post-vent | 100 | V3b, LO1, LO2, TO1, IPS2 | 26 |

### ECoG recordings

#### NYU

Stimuli were shown on a 15 in. MacBook Pro laptop. The laptop was placed 50 cm from the participant’s eyes at chest level. Screen resolution was 1280 × 800 pixels (33 × 21 cm). Prior to the start of the experiment, the screen luminance was linearized using a lookup table based on spectrophotometer measurements (Cambridge Research Systems). Recordings were made using one of two amplifier types: NicoletOne amplifier (Natus Neurologics, Middleton, WI), bandpass filtered from 0.16-250 Hz and digitized at 512 Hz, and Neuroworks Quantum Amplifier (Natus Biomedical, Appleton, WI) recorded at 2048 Hz, bandpass filtered at 0.01- 682.67 Hz and then downsampled to 512 Hz. Stimulus onsets were recorded along with the ECoG data using an audio cable connecting the laptop and the ECoG amplifier. Behavioral responses were recorded using a Macintosh wired external numeric keypad that was placed in a comfortable position for the participant (usually their lap) and connected to the laptop through a USB port. Participants self-initiated the start of the next run by pushing a designated response button on the number pad.

#### UMCU

Stimuli were shown on a NEC MultiSync® E221N LCD monitor positioned 75 cm from the participant’s eyes. Screen resolution was 1920 x 1080 pixels (48 x 27cm). Stimulus onsets were recorded along with the ECoG data using a serial port that connected the laptop to the ECoG amplifier. As no spectrophotometer was available at the UMCU, screen luminance was linearized by reverting the built-in gamma table of the display device. Data was recorded using a MicroMed amplifier (MicroMed, Treviso, Italy) at 2048 Hz with a high-pass filter of 0.15Hz and low-pass filter of 500Hz. Responses were recorded with a custom-made response pad.

#### Stimuli

All code used for generating the stimuli and for experimental stimulus presentation can be found at https://github.com/BAIRR01/BAIR_stimuli and https://github.com/BAIRR01/vistadisp.

Visual stimuli for the purpose of estimating changes in neural temporal dynamics were generated in Matlab 2018b. Stimuli were shown within a circular aperture with a radius of 8.3 degrees of visual angle using Psychtoolbox-3 (https://psychtoolbox.org/) and were presented at a frame rate of 60 Hz. Custom code was developed in order to equalize visual stimulation across the two recording sites as much as possible; for example, all stimuli were constructed at high resolution (2000×2000 pixels) and subsequently downsampled in a site-specific manner such that the stimulus was displayed at the same visual angle at both recording sites (see *stimMakeSpatiotemporalExperiment.m* and *bairExperimentSpecs.m*). Stimuli consisted of grayscale bandpass noise patterns that were created following procedures outlined in Kay et al., (2013). Briefly, the pattern stimuli were created by low-pass filtering white noise, thresholding the result, performing edge detection, inverting the image polarity such that the edges are black, and applying a band-pass filter centered at 3 cycles per degree (see *createPatternStimulus.m*). All stimuli were presented within a circular aperture; the remainder of the display was filled with neutral gray (**Fig. 1*A*)**. The spatial pattern we used has been shown to effectively elicit responses in most retinotopic areas (Kay et al., 2013b; Zhou et al., 2018).

**Figure 1:**
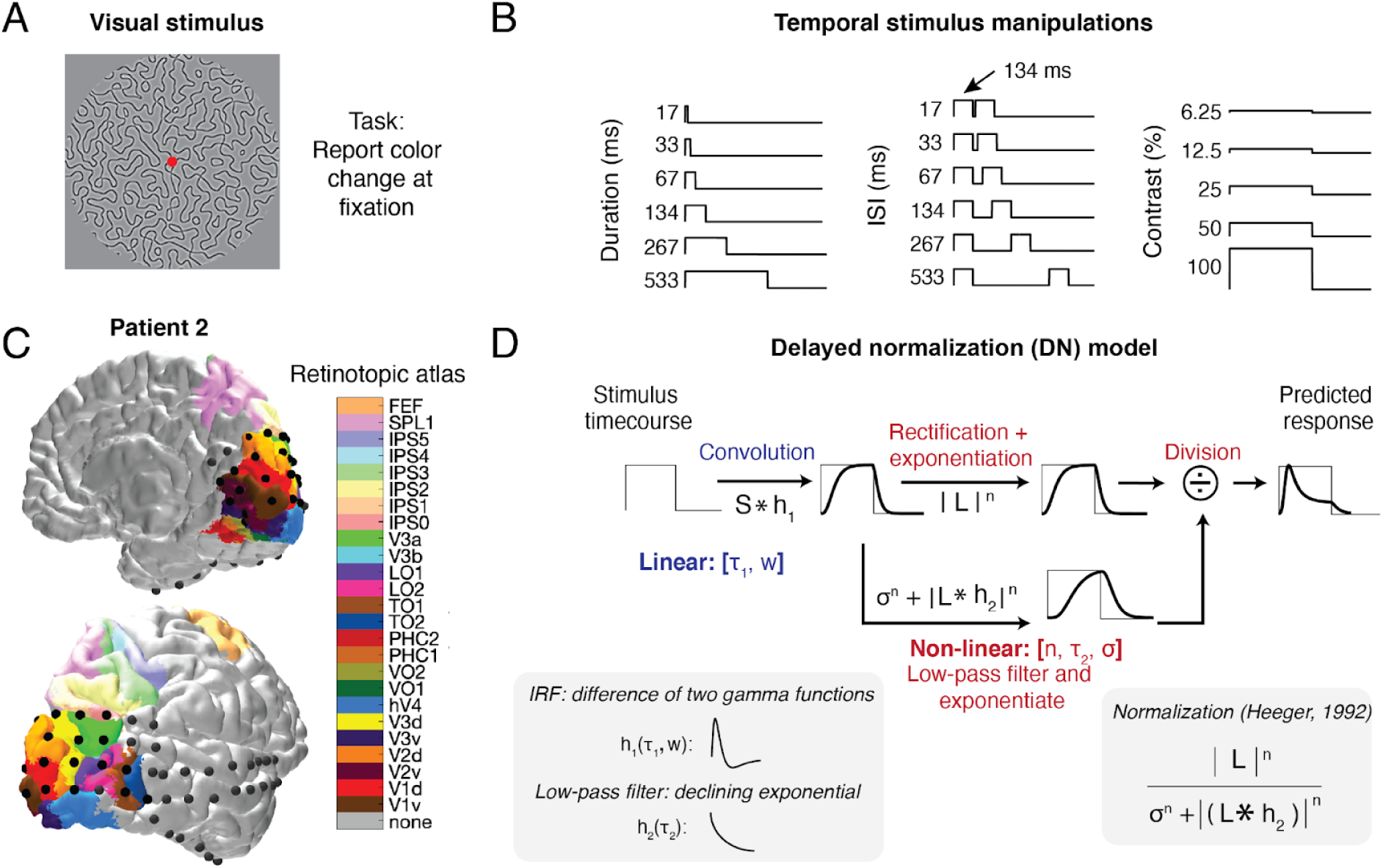
Experimental design, ECoG data and delayed normalization model. **A)** Participants were shown large (16°) grayscale pattern stimuli consisting of connected curved line segments. Unique individual stimuli were used for each stimulus condition. **B**) Stimulus time courses were varied in three different ways. For duration trials, a single full contrast stimulus was presented for different durations ranging between 17-533 ms. For repetition trials, a full contrast stimulus was presented twice for 134 ms, with an inter-stimulus interval (ISI) ranging between 17-533 ms. For contrast trials, a single stimulus was presented at a fixed duration (500 ms) with levels of contrast ranging between 6.25% to 100%. **C)** Electrode positions from an example participant overlaid on a pial surface reconstruction with predicted visual area locations. In this visualization, a surface node in the pial mesh was assigned a color if it had a non-zero probability of being on a visual region according to the full probability map from (Wang et al., 2015). If the electrode had a non-zero probability of being in multiple regions, the region with the highest probability was assigned. High-resolution images of each participant’s electrode positions and atlas projections are provided in **Extended Data Figure 1-1** and **Extended Data Figure 1-2**; interactive 3D figures can be generated using *tde_mkFigure1_1.m* in https://github.com/irisgroen/temporalECoG**. D)** Visual depiction of the delayed normalization (DN) model, first presented in Zhou et al., (2019). This linear-non-linear gain model takes a stimulus time course as input and produces a predicted response time course as output, by applying both linear and non-linear computations. The initial, linear part consists of convolution with an impulse response function (IRF; see lower left inset), parameterized as a difference of two gamma functions, with free parameters 𝜏_1_ (time constant) and *w* (weight of negative to positive gamma). This is followed by a non-linear part consisting of rectification, exponentiation with exponent *n*, and division by a semi-saturation contrast 𝜎, plus a delayed copy of the input that is also rectified and exponentiated. The delay is implemented as a convolution of the linear response with an exponentially decaying function of time constant 𝜏_2_ (see lower left inset). This model form constitutes a temporal implementation of divisive normalization, as first proposed by Heeger (1992) (see lower right inset). Fitted model parameters across electrodes in each visual area are provided in **Extended Data Figure 1-3.**

After generating independent pattern stimuli for each trial, stimuli were assigned to one of three different stimulus conditions (**Fig. 1*B*)**: duration, inter-stimulus-interval (ISI) or contrast (*stimMakeSpatiotemporalExperiment.m*). Duration and ISI trials were generated using the same parameters as in Zhou et al., (2018). For duration trials, a single pattern stimulus was shown at full contrast for one of 6 durations that ranged between 17-533 ms. For ISI trials, a single full contrast pattern stimulus was shown for a fixed duration (134 ms) twice in a row, with one of 6 inter-stimulus intervals whose duration varied between 17-533 ms. Duration and ISIs were powers of 2 times the monitor dwell time (1/60s). For contrast trials, contrast varied from 6.25% to 100% in 5 steps of powers of 2. Contrast trials were shown at a fixed duration of 500 ms. Together, all trial-types amounted to 17 stimulus conditions (6 duration, 6 ISIs, and 5 contrasts). The stimulus durations, contrast and inter-stimulus intervals were chosen to span a large dynamic range based on pilot data in V1-V3 in both fMRI and ECoG.

### Experimental design and statistical analysis

#### Experimental procedure

Duration, repetition, and contrast conditions were divided across two runtypes: one containing the duration- and ISI-varying trials, and one containing the contrast-varying trials as well as an additional set of grating and density-varying stimuli that were not analyzed for the purpose of the current paper. Trials were presented with an inter-trial-interval (ITI) that was randomly picked from a uniform distribution varying between 1.25 and 1.75 seconds. For each runtype, two unique stimulus sequences were created containing randomly ordered trials and ITIs. With the exception of participants 3 and 4 (who performed an earlier version of the experiment), all participants were presented with the same stimulus sequences. Each run contained 36 stimuli with 3 instances per stimulus condition and lasted about 60 seconds. Screen flip times, as measured by Psychtoolbox, were saved during stimulus presentation and compared to the requested timings to ascertain that stimuli were presented with high temporal accuracy (maximum estimated accumulated timing error per run <8ms; maximum deviation of stimulus duration <2 ms).

For all runtypes, participants were instructed to fixate on a cross located in the center of the screen and press a button every time the cross changed color (from green to red or red to green). Fixation cross color changes were created independently from the stimulus sequence and occurred at randomly chosen intervals ranging between 1 and 5 s. For each task, participants completed each unique run at least once. Runs were then repeated several times within the same experimental session or additional sessions recorded on different days. The amount of data collected for each patient is provided in **Table 1**. The experimenter stayed in the room during the experiment. Patients were encouraged to take short breaks between runs.

### ECoG data analysis

#### Data preprocessing

Data were read into Matlab 2020b using the Fieldtrip Toolbox (Oostenveld et al., 2011) and preprocessed with custom scripts available at https://github.com/WinawerLab/ECoG_utils. Raw data traces obtained in each recording session were visually inspected for spiking, drift or other artifacts. Electrodes that showed large artifacts or showed epileptic activity were marked as bad and excluded from analysis. Data were then separated into individual runs and formatted to conform to the ieeg-BIDS format (Holdgraf et al., 2019). Data for each run were re-referenced to the common average across electrodes for that run, whereby a separate common average was calculated per electrode group (e.g., one for grid and one for strip electrodes, see *bidsEcogRereference.m*). Next, a time-varying broadband estimate was computed for each run in the following way (see *bidsEcogBroadband.m*). First, the voltage traces were band-pass filtered using a Butterworth filter (passband ripples < 3 dB, stopband attenuation 60 dB) for 10 Hz-wide bands that ranged between 50-200 Hz. Bands that included frequencies that were expected to carry line noise and their harmonics were excluded (for NYU: bands 60, 120 and 180 Hz; for UMCU, 50, 100 and 150 Hz). The power envelope of each band-pass filtered voltage time course was calculated as the square of the time course’s Hilbert transform. The resulting envelopes were then averaged across bands by taking the geometric mean (see *ecog_extractBroadband.m*). Unlike the arithmetic mean, taking the geometric mean ensures that the resulting average is not biased towards the lower frequencies. The re-referenced voltage and broadband traces for each run were written to BIDS derivatives directories.

#### Electrode localization

Intracranial electrode arrays from NYU participants were localized based on pre- and post-implantation structural MRI images (Yang et al., 2012). Electrodes from UMCU participants were localized from the postoperative CT scan and co-registered to the preoperative MRI (Hermes et al., 2010). Electrode coordinates were computed in native T1 space and visualized onto pial surface reconstructions of the T1 scans generated using Freesurfer (http://freesurfer.net). Visual maps of striate and extrastriate cortex were generated for each individual participant based on the preoperative anatomical MRI scan by aligning the surface topology with two atlases of retinotopic organization: an anatomically defined atlas (Benson et al., 2014; Benson and Winawer, 2018) and a probabilistically defined atlas derived from a retinotopic fMRI mapping dataset (Wang et al., 2015); **Fig. 1*C***. Using the alignment of the participant’s cortical surface to freesurfer’s fsaverage subject, atlas labels defined on the fsaverage were interpolated onto their cortical surface via nearest-neighbour interpolation.

Electrodes were then matched to both the anatomical and the probabilistic atlases using the following procedure (see *bidsEcogMatchElectrodesToAtlas.m*): For each electrode, the distance to all the nodes in the freesurfer pial surface mesh were calculated, and the node with the smallest distance was determined to be the matching node. The atlas value for the matching node was then used to assign the electrode to one of the following visual areas in the anatomical atlas (henceforth referred to as the Benson atlas): V1, V2, V3, hV4, VO1, VO2, LO1, LO2, TO1, TO2, V3a, V3b, or none; and to assign it a probability of belonging to each of the following visual areas in the probabilistic atlas (henceforth referred to as the Wang atlas)): V1v, V1d, V2v, V2d, V3v, V3d, hV4, VO1, VO2, PHC1, PHC2, TO2, TO1, LO2, LO1, V3b, V3a, IPS0, IPS1, IPS2, IPS3, IPS4, IPS5, SPL1, FEF, or none.

#### Data selection

The preprocessed data were analyzed for the purposes of this paper using custom Matlab code available at https://github.com/irisgroen/temporalECoG. First, a dataset was created by reading in the voltage and broadband traces of each run from each participant from the corresponding BIDS derivatives folders (*tde_getData.m*). Only electrodes that had a match with one of the visual areas from the anatomical atlas (Benson and Winawer, 2018) or one the visual areas in the maximum probabilistic atlas from (Wang et al., 2015) were selected for preprocessing. Only (high-density) grid or strip electrodes were included (i.e. depth electrodes were excluded).

In order to combine the UMCU and NYU participants in a single dataset, the UMCU data was resampled at 512 Hz. Broadband and voltage traces from each run were subsequently segmented into individual epochs by extracting the neural time courses between [-0.1 and 1.2] seconds relative to stimulus onset. Visual inspection of the data indicated an obvious delay in response onset for the UMCU participants relative to the NYU participants. The cause of the delay could not be tracked down but it was clearly artefactual. To correct the delay, UMCU data were aligned to the NYU data based on a cross-correlation of the average event-related potential (ERP) across all stimulus conditions from V1 electrodes from three participants (1 UMCU, 2 NYU). The delay in stimulus presentation was estimated to be 72ms (95% CI 63 - 85 ms by bootstrapping across a total of 18 electrode pairs), and stimulus onsets of all trials from the UMCU participants were shifted according to this average delay (see *s_determineOnsetShiftUMCUvsNYU.m*)

Voltage epochs were baseline-corrected by subtracting the average pre-stimulus amplitude within each trial. Broadband epochs were converted to percent signal change by pointwise dividing and subtracting the average prestimulus baseline (i.e., average broadband power between -100 and 0 ms relative to stimulus onset) across all epochs within each run. We then performed two consecutive data trimming steps: 1) epoch selection and 2) electrode selection (see *tde_selectData.m)*.

Epoch selection was performed using both the voltage and broadband epochs. Epochs were automatically rejected according to two criteria (see *ecog_selectEpochs.m*): 1) a difference in voltage between consecutive samples larger than 200 mV anywhere in the epoch; 2) the maximum broadband amplitude in the epoch *outside* the time window [0.05 0.85] relative to stimulus presentation exceeds the average of the maximum response across trials *inside* the stimulus presentation window by 3 standard deviations. Across participants, on average 2.5% of epochs were rejected (sd = 0.6%, min = 1.7%, max = 3.2%).

Electrode selection was performed based on the broadband epochs and the electrode locations. To be included in data analysis, electrodes had to meet one criterion (see *ecog_selectElectrodes.m*): an above-threshold correlation between two independent halves of the data (split-half reliability). To compute this measure for each electrode, all epochs for a given trial type were randomly assigned to one of two splits. Within each half, epochs of the same trial-type were averaged and then concatenated, and the coefficient of determination (R^2^) was calculated between the two concatenated time series. Electrodes with a low split-half reliability (R^2^ < 0.22) were excluded.

For three participants, this electrode selection procedure resulted in all their electrodes getting rejected from analysis. For the 8 remaining participants, 70.5% of electrodes on average were rejected (sd = 14.5%, min = 50%, max = 92%; see **Table 1** for subject-specific data).

#### Data summary

The data preprocessing, electrode localization and data selection procedures outlined above resulted in a total of 98 electrodes with robust visual responses covering visual areas V1, V2, V3, V3a, V3b, hV4, LO1, LO2, TO1, TO2, IPS0-4 (see **Table 2** for an overview of electrodes per area). The broadband epochs of selected electrodes were averaged across all trials within a stimulus condition, resulting in 17 response time courses per electrode that constituted the data for computational model fitting (see below).

**Table 2:** Overview of electrodes and patients per visual area. Columns refer to the following: *Visual area:* Name of retinotopic map. Ventral and dorsal maps of early visual areas (V1-3) were combined. Areas TO1 and TO2 were combined into a single area TO, and IPS0-4 were combined into a single area IPS (see Materials and Methods). *Number of electrodes*: Number of electrodes with non-zero overlap with each retinotopic area, according to the full probability retinotopic atlas by Wang et al., (2014). Note that the sum is larger than the total number of electrodes included in the dataset (n=98), because electrodes can probabilistically map onto several areas. *Median number of electrodes per bootstrap*: For a given bootstrap, electrodes are sampled with replacement and probabilistically assigned to one visual area at a time. Note that the sum is smaller than the total number of electrodes included in the dataset because some electrodes had a non-zero probability of overlapping with hV4, which was excluded from this report (see Methods), but was still included in the area assignments. *Contributing patients*: List of patients that contribute electrodes for each visual area.

| Visual area | Number of electrodes | Median number of electrodes per bootstrap | Contributing patients |
| --- | --- | --- | --- |
| V1 | 12 | 7 | patient 2, 3, 6, 7, 10 |
| V2 | 37 | 12 | patient 2, 3, 4, 5, 6, 7, 10 |
| V3 | 56 | 16 | patient 2, 3, 4, 5, 6, 7, 10, 11 |
| V3a | 35 | 4 | patient 3, 4, 7, 10, 11 |
| V3b | 50 | 17 | patient 3, 4, 7, 10, 11 |
| LO1 | 73 | 19 | patient 3, 4, 5, 7, 10, 11 |
| LO2 | 34 | 9 | patient 3, 10, 11 |
| TO | 12 | 3 | patient 3, 10, 11 |
| IPS | 23 | 4 | patient 7, 10, 11 |

#### Probabilistic electrode assignment

When assigning individual electrodes to visual areas, we used a bootstrapping procedure that took into account the probability of each electrode overlapping with a visual retinotopic region. In this procedure, we repeatedly (for n bootstraps) assigned electrodes to visual areas in accordance with each electrode’s matched node’s probability distribution across areas determined by the Wang probability atlas (Wang et al., 2015), resulting in a distribution of bootstrapped electrodes for each visual area. From these distributions we then computed summary statistics (median or mean) and confidence intervals across neural responses or model predictions for each visual area. Before conducting the repeated assignments, we rescaled the probability values in the atlas to exclude the ‘none’ probabilities, which ensured that each electrode would always get assigned to an area (e.g. if an electrode had 60% probability of belonging to V3b, 20% probability of belonging to V3a, and 20% probability of belonging to ‘none’, it would get assigned to V3b 75% of the time and to V3a 25% of the time). To guarantee robust estimates, we only report results from areas that had at least 10 electrodes with a non-zero probability of being positioned on that area. This resulted in the exclusion of area hV4. We additionally grouped together TO1 and TO2 into a single area TO.

### Computational modeling

#### Model fitting: temporal dynamics

We fit models of temporal dynamics to the 17 broadband time courses for each individual electrode (see *tde_fitModel.m*). Each model takes stimulus time courses as input and predicts neural response time courses as output. For analyses focusing on describing different temporal phenomena in V1 (Figures 2-4), and comparisons of model performance (Figure 7), models were fit separately to individual electrodes, and parameters or metrics derived from these fits were then averaged within visual areas using the probabilistic assignment bootstrapping procedure described above. For analyses that characterized changes in temporal dynamics along the visual hierarchy (Figures 8-10), we instead conducted repeated (n=1000) fits to the average time courses across electrodes within an area, whereby for each fit the average time course per area was computed after performing the probabilistic assignment, yielding a slightly different average and thus different model fit each time. While results and conclusions were not qualitatively different, we found that in higher visual areas, this averaging-then-fitting procedure yielded slightly more robust estimates compared to fitting individual electrodes and subsequently averaging them. For analyses illustrating similarities in contrast and adaptation time courses (Figures 5-6), model predictions were generated after electrodes were first averaged probabilistically within V1.

Models were fitted using Bayesian adaptive direct search (BADS; Acerbi and Ma, 2017). BADS alternates between a series of fast, local Bayesian optimization steps and a systematic, slower exploration of a mesh grid (see https://github.com/lacerbi/bads for code and further explanation of the algorithm). Using BADS resulted in slightly more robust fits to the data compared to two built-in Matlab alternatives *fmincon* and *lsqnonlin*; however, results and conclusions did not change qualitatively when using these optimizers, and we implemented all three options in our fitting code (*tde_fitModel.m*). Model performance was evaluated using the cross-validated coefficient of determination (R2) computed via 17-fold leave-one-out cross-validation whereby each of the stimulus conditions was once left out of the fitting procedure and then predicted based on the parameters estimated from the remaining 16 stimulus conditions. Model R2 was computed by averaging across the R2 values for the 17 left out stimuli. Model parameter values and summary parameters (see below) were estimated based on fits to the full dataset.

#### Computational models of temporal dynamics

The DN model and its reduced versions (see below) are implemented as standalone functions in https://github.com/irisgroen/temporalECoG, folder ’temporal_models’. Each model is paired with a JSON metadata file containing parameter descriptions, starting points and upper and lower bounds that were used when fitting the model. As detailed below, models differ in the number and types of parameters fitted. In addition to the model-specific parameters, two nuisance parameters were fitted for all models: *shift* (delay in response onset relative to stimulus onset) and *scale* (gain of the response), to take into account differences between electrodes or visual areas in response onset latency and overall response magnitude (i.e. responses in V1 are much higher than in later visual areas). We tested the following models:

#### Delayed normalization model

Our main model of interest was the delayed normalization (DN; **Figure 1D**) model previously described in Zhou et al., (2019). The core idea of the model is that the stimulus drive is divisively normalized by delayed population activity (here, delayed via a low pass filter). There is a long history of implementing models with a delay in the normalization pool, including (Heeger, 1992, 1993; Mikaelian and Simoncelli, 2001; Tsai et al., 2012; Sinz and Bethge, 2013); see Zhou et al., 2019, Figure 7, for more elaborate discussion on the relation of the DN model with other implementations of delayed gain control mechanisms. Because prior model implementations were either conceptual, meaning not fit to data (e.g., Heeger, 1992) or fit to specific types of data (eg., Zhou et al., 2019), it was necessary to implement a specific formulation that could be applied to a wide variety of ECoG data, as we have here. Zhou et al., (2019) demonstrated that the DN model was able to predict compressive temporal summation (in ECoG data to stimuli of a single duration), adaptation (in fMRI responses to stimuli of varying durations and interstimulus-intervals) and contrast dynamics (in non-human primate MUA recordings from V1). Here, we tested whether the same model could predict all three of these phenomena *simultaneously* in ECoG data. We largely re-use Zhou’s 2019 implementation, which included a parameterized impulse response function, exponentiation of the driven activity, and an exponential decay filter applied to the normalization pool. The DN model is defined by an LNG structure (Linear, Nonlinear, Gain control) and is described by the following set of equations:

First, a linear response (R*_L_*) is computed by convolving an input stimulus time course *S* of length *t* (dimensions 1x*t*, with values ranging between 0 and 1) with an impulse response function (IRF) *h_1_*, consisting of a weighted difference of two gamma functions:

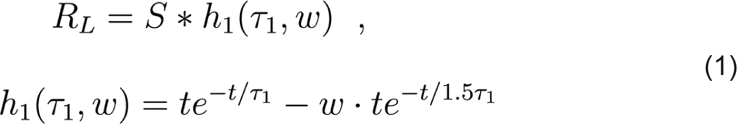

in which *t* is time, *w* the weight on the second (negative) gamma function, and 𝜏_1_ is a time constant determining the shape of the IRF. Symbols in parentheses are parameters. *R_L_*, *S*, and *h_1_* are functions of time; we omit *t* for simplicity. Note that each individual gamma function here is a simplification of the gamma function provided in Boynton et al., (1996); their equation 3, whereby the phase delay of the IRF is fixed at value 2 (as in Zhou et al., 2019). Moreover, as in Zhou et al., (2019), we assumed the peak timing of the second (negative) gamma function to be 1.5 times the first (positive) one, leaving only 𝜏_1_ and *w* as the free parameters of the linear step. This version of the DN model (*DN.m*) was used for all analyses with the exception of the model reduction analysis (Figure 7), where we compare this model to models with a more flexible IRF, which has two additional free parameters: one for the phase delay and one for the time constant of the negative gamma function (see next section).

The linear response (R*_L_*) is then converted into a non-linear response R*_LN_* by applying a full-wave rectification and an exponentiation with exponent *n*, which is fitted:

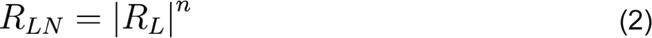

The final step is divisive normalization of the non-linear response R*_LN_* with a low-pass filtered version of R*_L_* that is also rectified and exponentiated to the same *n*. In addition, an exponentiated semi-saturation constant 𝜎 is added to the denominator:

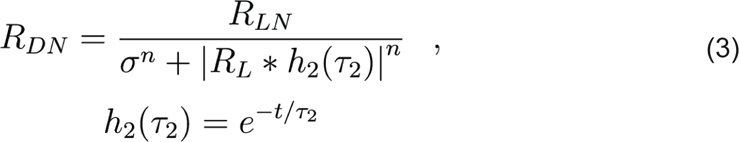

The low-pass filtering here is achieved by means of convolution with an exponential decay function *h_2_* with time constant 𝜏_2_.

As noted in **Fig. 1*D*,** equation 3 has the form of canonical divisive normalization (Carandini and Heeger, 2012), in which a numerator (reflecting the rectified and exponentiated linear input) is divided by a denominator consisting of a normalization pool and a semi-saturation constant, which each are also raised to the same exponent *n* as the numerator. Here, the normalization pool consists of a delayed version of the numerator, which means that the input drive is essentially normalized by a delayed version of itself. This yields an output time course that is characterized by a transient response rise followed by a decay to a sustained response level.

To summarize, for the main version of the DN model used throughout, five parameters were fitted in total: 𝜏_1_ (time constant of the IRF), *w* (weight of the negative and positive IRFs), *n* (exponent), 𝜎 (semi-saturation constant), and 𝜏_2_ (time constant of the exponential decay). Model parameters and bounds were initialized using values used in Zhou et al., (2019).

#### Delayed normalization model, fully parameterized and deconstructed

In addition to the original model proposed by Zhou et al., (2019), we tested reduced versions of that same model, in which non-linear computations were consecutively added on top of the purely linear first step of the model. The motivation for including this model construction analysis was to investigate how much each step contributed to the overall predictive performance of the model. As explained above, we additionally adapted the model such that IRF used for linear convolutions was less constrained, allowing the time constant of the negative gamma function and the phase delay of both gamma functions to be fitted in optimization, such that it could capture as much variance in the data as possible with a maximally flexible IRF (total number of fitted parameters = 4: 𝜏_1pos_, 𝜏_1neg_, phase delay *r*, and *w*; see *LINEAR.m*). We then added full-wave rectification (*LINEAR+RECTF.m*; same parameters as *LINEAR.m*), exponentiation (*LINEAR+RECTF+EXP.m;* adding parameter *n*), normalization without delay (*LINEAR+RECTF+EXP+NORM.m;* adding parameter 𝜎) and finally, normalization with delay (*LINEAR+RECTF+EXP+NORM+DELAY.m*; adding parameter 𝜏_2_), which is equivalent to the DN model (*DN.m*), except that the IRF is less constrained. Model parameters and bounds were initialized using the same values as for the DN model and can be found in the JSON meta-data of the fitting code in https://github.com/irisgroen/temporalECoG.

#### Summary metrics

To characterize the temporal dynamics of the broadband time courses, we computed several summary metrics on both data and model predictions (see *tde_computeDerivedParams.m*):

*Time-to-peak*: The time interval between stimulus onset and the maximum (peak) of the response time course (in seconds). We computed this based on responses or model predictions for the longest duration, full contrast stimulus (533 ms), which is the condition of maximal stimulation and thus the conditions where the response is overall expected to be greatest..

*Full-width half max (Fwhm)*: The difference between the time-point (relative to stimulus onset) at which the response has risen to half of the maximum, and the time-point at which the response has decayed again to the same half-way point (in seconds). This metric was computed based on neural response or model prediction time courses to short duration stimuli (e.g. 17 ms), which have the least amount of sustained level activation and therefore the most symmetric response shapes.

*Ratio sustained/transient*: Response magnitude at the sustained level divided by the transient response magnitude (maximum/peak). For estimates of the sustained level, we took the response level at stimulus offset for the longest duration stimulus in the dataset (533 ms). To achieve robust estimates of the offset, data and model predictions were smoothed using a moving average filter with a span of 150 data samples (∼290) ms (see *tde_mkFigure4.m*). The moving average is symmetric around each data point and hence does not systematically shift the timing of the peak.

*Recovery from adaptation*: The relative magnitude of response of the second stimulus relative to the first stimulus in the repeated stimuli conditions. The goal of this procedure is to estimate which part of the neural time course reflects the continuing activation to the first stimulus and which part reflects new neural activity elicited by the second stimulus. For each electrode, we compute an estimate of the response to the first stimulus by averaging together 1) the time course of an actual 134 ms stimulus presented in isolation (the fourth duration-varying condition), which includes all time points in entire epoch, and 2) each of the ISI-varying conditions, but only including time-points up to the onset of the second stimulus (see *tde_computeISIrecovery.m)*. The reason for additionally including these ‘partial time courses’ instead of using the 134 ms duration stimulus only, is that it gives us a more robust estimate of the response to the first stimulus (e.g. of the transient peak). Importantly, the estimated response to the first stimulus never includes data points after a second stimulus was presented. We then subtracted this average time course from each of the ISI-varying stimulus responses, yielding an estimate of the response of the second stimulus ’corrected for’ the response to the first stimulus. We then computed relative recovery by taking the maximum (peak) of the corrected second stimulus time courses and comparing it with the peak response of the average first stimulus estimate. Using the summed response across all time points in the time course for which a stimulus was present rather than the peak of the time course yielded qualitatively similar results.

*C50*: Percentage contrast for which response magnitude reaches 50% of the response to a 100% contrast stimulus. For both the neural data and the model predictions, this point was estimated by fitting a Naka-Rushton function (Naka and Rushton, 1966; Albrecht and Hamilton, 1982) to the maxima of the response time courses for each of the 5 contrast levels we measured. This equation takes the following form:

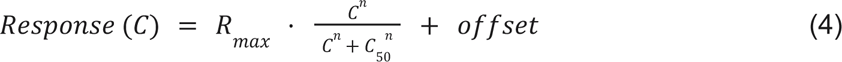

Where *C* is the stimulus contrast level, R*_max_*is the maximum response amplitude, *C50* the contrast at which the curve reaches half height, and exponent *n* controls the steepness of the sigmoid (see *fitNakaRushton.m*). Taking the sum rather than the peak of the time course as the response per contrast level yielded qualitatively similar results.

### Code and data availability statement

All code used for the purpose of this paper can be found at the GitHub repositories mentioned above. The ieeg-BIDS-formatted data, derivatives, the processed dataset and model fits are available on OpenNeuro (<u>doi:10.18112/openneuro.ds004126.v1.0.0</u>)

## Results

We first examined how each of our three stimulus manipulations (duration, repetition and contrast) affected the temporal dynamics of ECoG broadband time courses in area V1.

### Neural responses in human V1 exhibit subadditive temporal summation

First, systematic variations of stimulus duration demonstrated the presence of *subadditive temporal summation* in neural responses in V1 (**Fig. 2**). When a stimulus doubles in duration, its resulting neural response is reduced relative to the linear prediction, i.e., a sum of the responses to the shorter stimulus and a time-shifted copy of that response (**Fig. 2*A***). This demonstrates that the additional visual exposure resulting from a longer presentation duration does not accumulate linearly. Comparison of summed neural responses across all stimulus durations indeed shows evidence of subadditive summation; throughout the 30-fold range measured in our experiment (17 ms to 533 ms), the obtained responses deviate systematically from the linear prediction, with responses to longer stimuli much smaller than the linear prediction extrapolated from the briefest stimulus (**Fig. 2*B*)**.

**Figure 2:**
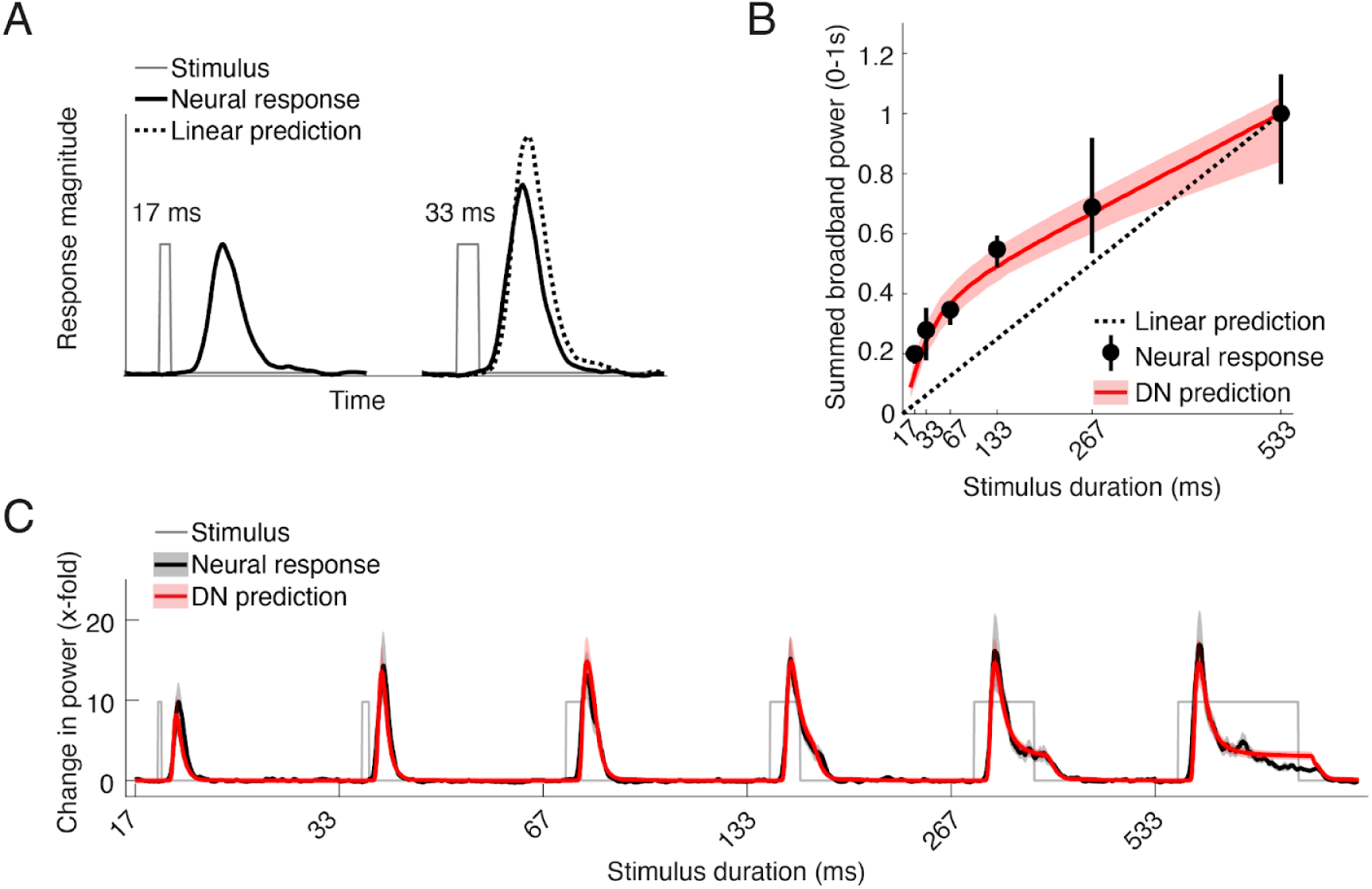
Sub-additive temporal summation in neural responses in human V1. **A)** Compressive temporal summation in neural responses illustrated by two stimulus time courses (17 and 33 ms durations, gray thin lines) and corresponding ECoG broadband time courses (black thick lines). Doubling stimulation duration does not result in a doubling of the response, as predicted by a linear prediction, computed as the sum of the shortest duration response and a shifted and scaled copy of that response (dotted line). **B**) Sum of broadband response time courses between 0-1 s after stimulus onset for each stimulus duration (black) and the predicted response by the DN model (red), normalized to the response to the longest duration stimulus (533 ms). Compressive temporal summation is evident as deviation of both data and model from the linear prediction extrapolated from the longest duration stimulus (dotted line). Data points indicate median across repeated samples of V1 electrodes (total n = 12; median n per bootstrap = 7; see **Table 2**), taking into account their probability of overlap with a retinotopic atlas (see Methods). Error bars and shaded regions reflect 68% confidence intervals across 10,000 bootstraps of electrode assignments. **C**) Average ECoG broadband time courses in V1 (black) for all stimulus durations measured (17-533 ms), along with the average prediction by the DN model (red). Shaded regions reflect 68% confidence intervals across 1000 bootstraps of electrode assignments. This figure can be reproduced with *tde_mkFigure2.m* in https://github.com/irisgroen/temporalECoG.

The reason for this can be understood by looking at the average response time courses themselves (**Fig. 2*C***). The largest response is the initial transient. For short duration stimuli, this is the only part of the response. At longer durations, there is also a lower-amplitude sustained response. Summed responses in **Fig. 2*B*** reflect the combination of both the transient and sustained level activity; at longer durations, the lower-amplitude sustained component comprises an increasingly large fraction of the response.

We find that this temporal subadditivity in ECoG broadband responses is well captured by a delayed normalization (DN) model (**Fig. 1*D***). The model accurately predicts the rapid increase in response at short durations and the slower increase at longer durations (**Fig. 2*B***). Moreover the model time courses accurately fit both the transient and the sustained levels of neural response across different stimulus durations (**Fig. 2*C***).

### Neural responses in human in V1 exhibit response suppression due to adaptation

Second, repetitions of stimuli with different inter-stimulus intervals (ISIs) demonstrated the presence of *adaptation* in neural responses in V1 (**Fig. 3**). Adaptation is evident as a reduction in stimulus-evoked response due to the presence of a preceding stimulus, and is most pronounced when the two stimuli are presented close together in time (**Fig. 3*A***). Adaptation effects are non-linear: while a linear model predicts no change in response magnitude due to preceding stimuli, the neural data reveal strong suppression for a stimulus that is presented in close temporal proximity to another stimulus, which gradually recovers as the ISI increases (**Fig. 3*B***). These adaptation effects are clearly visible in the response time courses to the separate stimuli (**Fig. 3*C***), where it can be seen that the response to the second stimulus achieves near-full recovery by the longest ISI tested (533 ms). Notably, these adaptation effects are well captured by the delayed normalization model, as indicated by the red lines in **Fig. 3*B-C***, although there are slight underpredictions for the response to the second stimulus at short ISIs.

**Figure 3:**
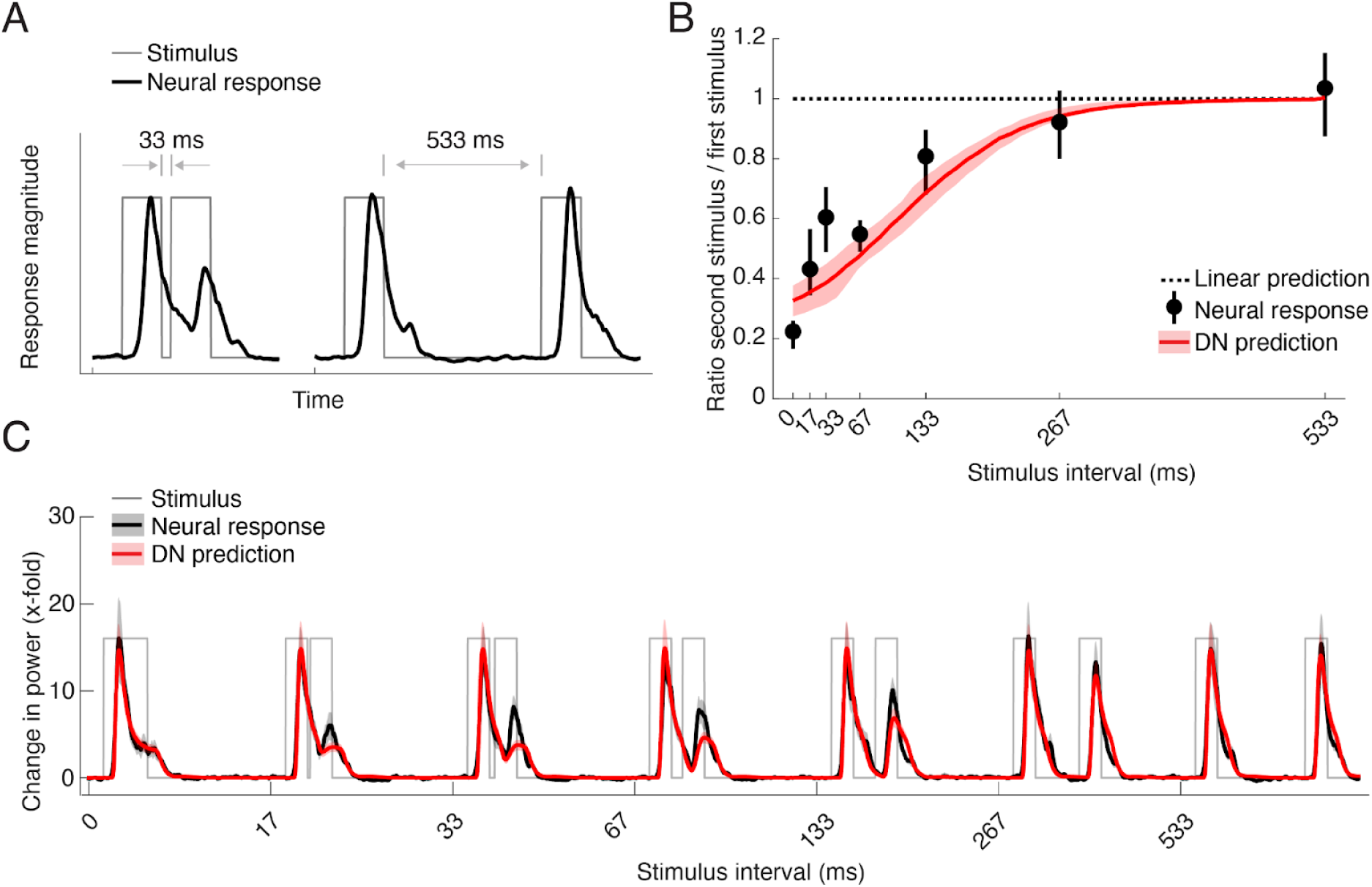
Adaptation of neuronal responses to stimulus repetition in human V1. **A)** Example of adaptation to a repeated stimulus, showing time courses for two stimuli of equal duration with a short (33 ms) and a long (533 ms) inter-stimulus interval (gray thin lines) and their corresponding ECoG broadband time courses (black thick lines). Short intervals lead to a strong suppression of the response to the second stimulus, and the response recovers with longer intervals. **B**) Recovery from adaptation for each condition, expressed as the difference in maximum response for the second stimulus compared to the maximum response of the first stimulus (black dots) along with the predicted recovery by the DN model (red line). Adaptation is evident from the deviation of both data and model from the linear prediction (dotted line). Data points indicate median across probabilistically assigned V1 electrodes (total n = 12; median n per bootstrap = 7; **Table 2**). Error bars and shaded regions indicate 68% confidence intervals across 10,000 bootstraps of electrode assignments. **C**) Average broadband time courses (black) in V1 for inter-stimulus intervals between 0-533 ms, along with predictions by the DN model (red), computed as in **Fig. 2*B***. This figure is produced using *tde_mkFigure3.m*.

### Neural responses in human in V1 exhibit slower dynamics with reduced contrast

Third, we observed *contrast-dependent temporal dynamics* of neural responses in V1, in spite of the fact that the temporal structure of the stimulus time course itself did not differ across the contrast conditions (each stimulus was presented at a single duration of 500 ms). Increasing stimulus contrast resulted not only in an increase in peak response magnitude (**Fig. 4*A***, left), but the response shape also changed in other ways independent of the peak magnitude, showing increased time-to-peak with lower contrast (phase delay) and a relatively less pronounced transient relative to the sustained level response (**Fig. 4*A***, right). This result differs from the descriptive model in Albrecht et al., (2002) in which the effect of contrast is modeled as a shift and scale in the response shape.

**Figure 4:**
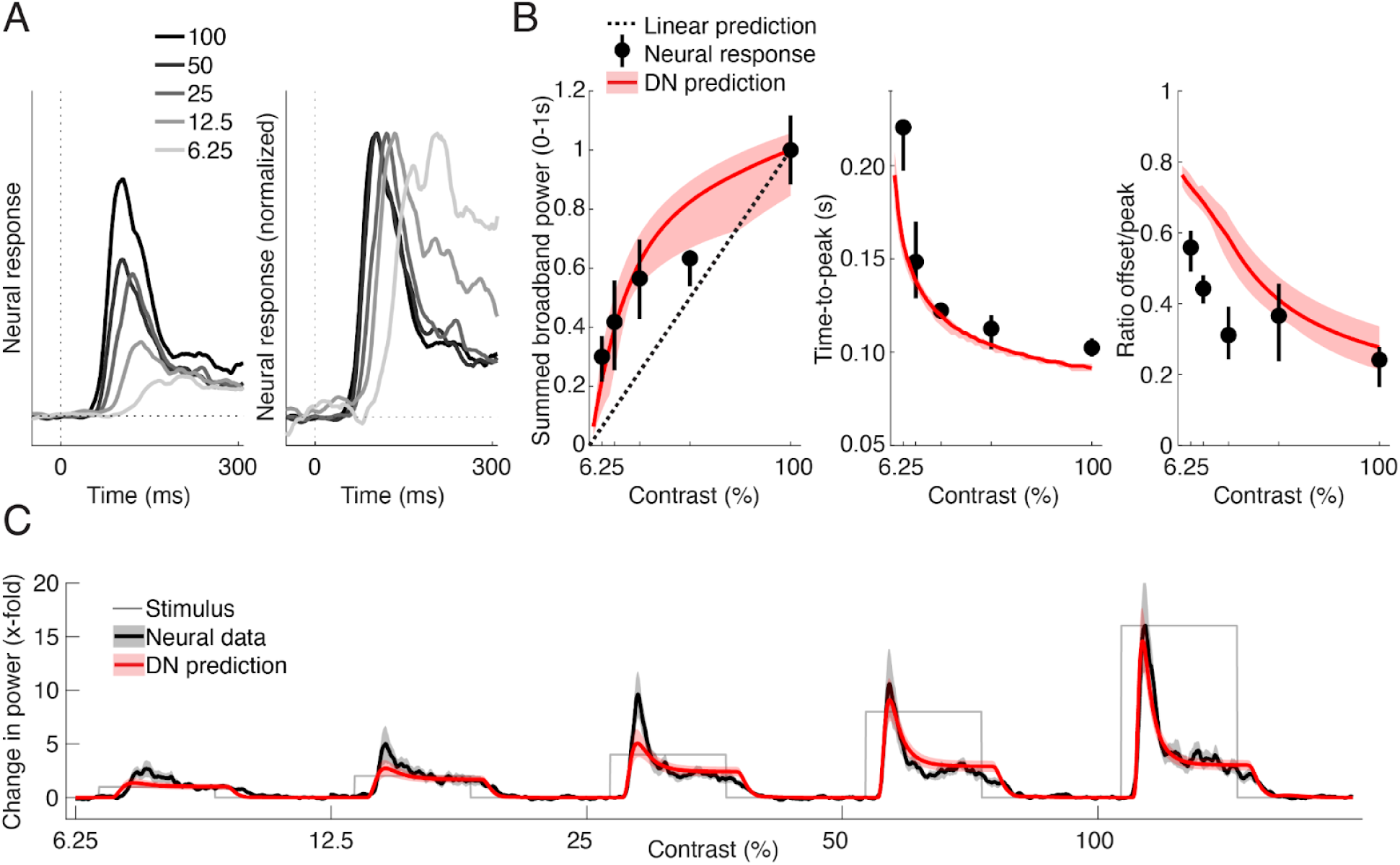
Contrast-dependent temporal dynamics of neuronal responses in human V1. **A)** Average ECoG broadband responses in V1 for different levels of contrast (6.25%-100%), superimposed (left) and superimposed and peak-normalized (right). Decreasing stimulus contrast results not only in a decrease in peak amplitude (left), but also a shift in peak latency and a decrease in the ratio of transient to sustained level of response (right). **B**) Summary statistics of contrast-dependent responses derived from individual electrodes (black dots) and DN model predictions (red line). Left: contrast response functions, measured as summed broadband power between 0.05-1s after stimulus onset. Middle: Time to peak. Right: Ratio of sustained response level (response magnitude at stimulus offset) to transient response level (maximum response level). Data points indicate median across probabilistically assigned V1 electrodes (total n = 12; median n per bootstrap = 7; **Table 2**). Error bars and shaded regions reflect 68% confidence intervals across 10,000 bootstraps of electrode assignments. **C**) Average ECoG broadband time courses in V1 (black) along with predictions by the DN model (red), computed as in **Fig. 2*B***. This figure is produced using *tde_mkFigure4.m*

We quantified these dynamics by computing three different summary statistics from each electrode’s response time course at each contrast level, which each again showed deviations from linearity (**Fig. 4*B***). Summed responses over time give rise to a contrast response function that progressively increases for higher contrasts (**Fig. 4*B***, left). Estimates of time-to-peak at each contrast level show that peak latency sharply decreases with increasing contrast (**Fig. 4*B***, middle). A comparison of the sustained level response (at stimulus offset) compared to the transient level (peak response) shows that the difference between these two measures decreases as contrast decreases, due to a less pronounced transient (**Fig. 4*B***, right). All three effects are again also qualitatively captured by the DN model, resulting in highly accurate model predictions across all contrast levels (**Fig. 4*C***). The model predictions are not perfect, however; they slightly underestimate the height of the transient part of the response for lower contrasts, resulting in elevated estimates of the sustained/transient response ratio (i.e., values closer to 1; **Fig. 4*B***). This slight model failure parallels the underprediction of the response to a second stimulus at short ISIs.

In sum, the V1 results show that there are several robust non-linear temporal dynamics in ECoG responses in human V1 that are all consistent with a time-dependent normalization, and thus well described by the DN model. We note that the model captures well-known nonlinear phenomena (e.g. contrast response function), as well as phenomena that are perhaps less well-characterized in population responses in human visual cortex, such as temporal summation and short-latency visual adaptation.

### Contrast and repetition effects lead to similar response reductions in V1

An advantage of testing many stimuli in the same experiments is that we can directly compare the effects of different stimulus manipulations. We showed nonlinearities with respect to contrast manipulations (**Fig. 4**) and with respect to stimulus repetition (**Fig. 3**). Here we ask how the two effects compare, and how the DN model simultaneously accounts for them both.

First, zooming into the broadband time courses contrast responses (same data as in **Fig. 4*A***) at the first 100 ms after stimulus onset clearly shows that lower stimulus contrast leads to more slowly rising responses, and to a change in response shape, with less distinction between transient and sustained responses (**Fig. 5**, top left). Interestingly, responses to repeated stimuli, after subtracting out the influence of the first stimulus (see Methods) have remarkably similar shapes (**Fig. 5**, top right). The similarity of the effects of both contrast and repetition on the response dynamics is predicted by the DN model (**Fig. 5**, bottom row). Therefore, both the data and model suggest the possibility that V1 responds to a recently viewed stimulus similarly to a novel but low-contrast stimulus. One difference between model prediction and data is that in the predictions, the responses for all stimuli start rising at the same time (but at different rates), whereas in the data, the onset latency appears to be delayed for low contrast and short repetitions.

**Figure 5:**
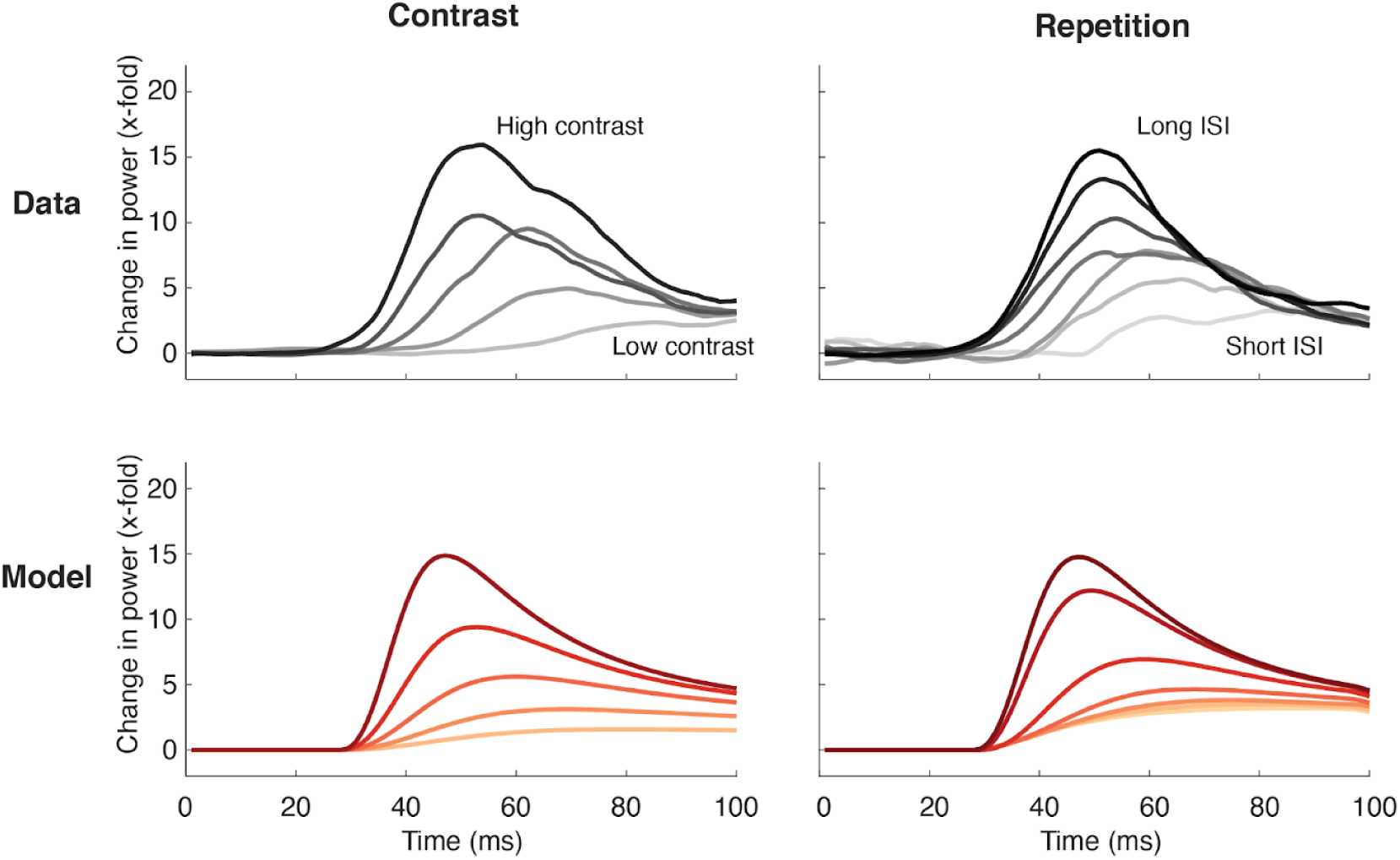
Comparison of contrast and repetition effects in ECoG data and DN model predictions. Top left: Average broadband time courses in V1 for the contrast-varying stimuli during the first 100 ms after stimulus onset. Darker lines indicate increasing contrast. Responses are stronger and rise faster with increasing contrast. Top right: Average broadband time course for the second stimulus in each repetition condition, after subtracting out the response to the first stimulus (see Methods). Darker lines indicate increasing ISI. Responses are stronger and rise faster with longer ISIs. Bottom row: DN model predictions derived using parameters from fitting to the average time courses in top row, for contrast-varying stimuli (left) and repeated stimuli (right). This figure is produced using *tde_mkFigure5.m*.

Inspecting the inner workings of the delayed normalization model helps understand why these responses look so similar and how the model accounts for the two effects simultaneously. In the model, the input drive is represented by the numerator, while the normalization pool is represented by the denominator (see **Fig. 1*D***). In **Figure 6**, these two components of the model are plotted separately for two different contrasts and two different ISIs. This shows that contrast and adaptation effects are both driven by differences in the normalization pool (denominator), but that for contrast, the effect can be attributed to the semi-saturation constant (i.e., left hand of denominator in **Fig. 1*D***), while for repetition, the effect can be attributed to the normalized input drive (i.e., the right hand of the denominator).

**Figure 6.**
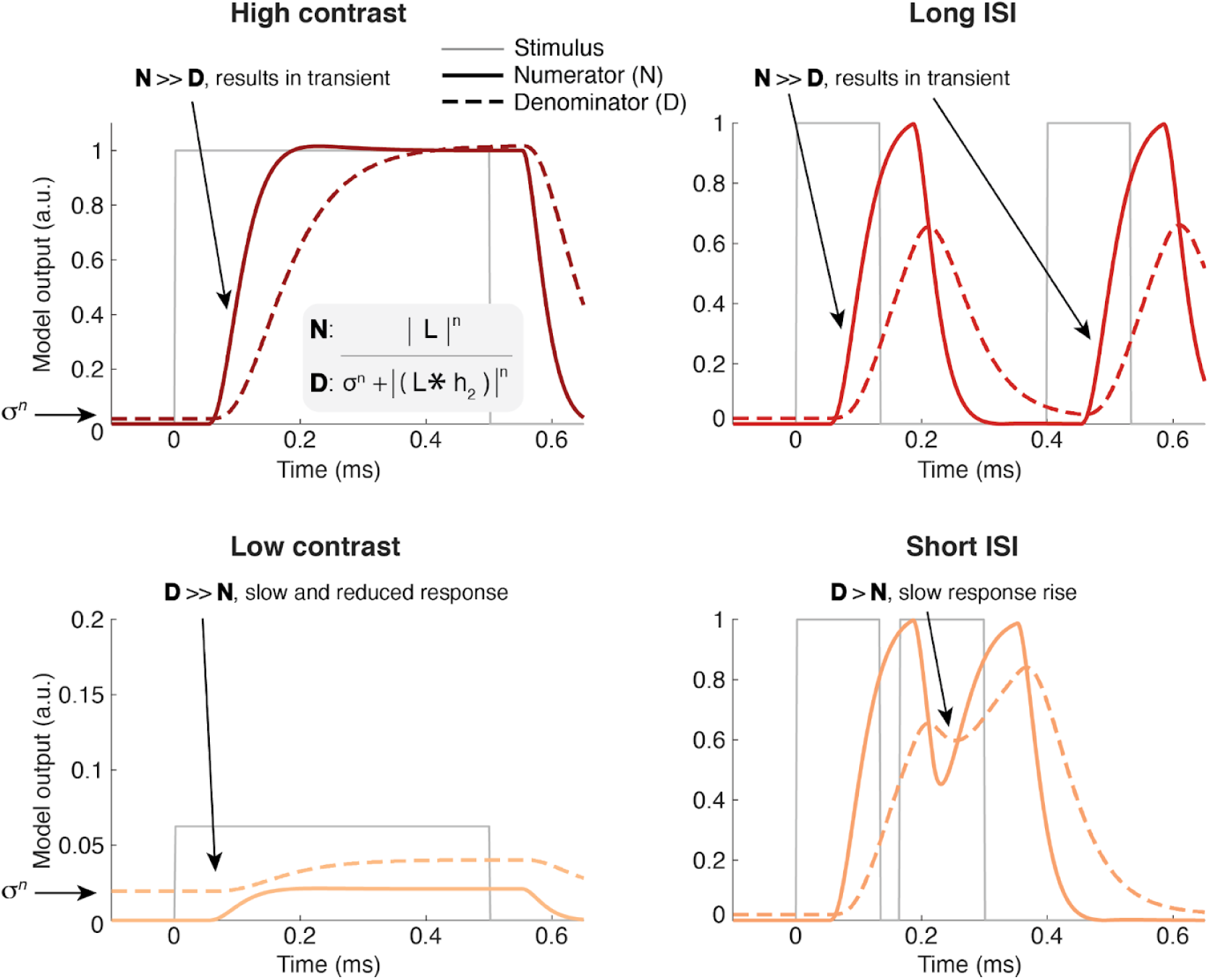
Illustration of how the DN model predicts similar effects of low-contrast and repetition. DN model numerator (solid lines) and denominator (dashed lines) for high contrast (top left), low contrast (bottom left), long ISI (top right) and short ISI (bottom right), for the model parameters fit to the average V1 response. (But note that no gain factor is applied, so the output units are relative to stimulus contrast.) With high contrast, the numerator quickly rises and thereby diminishes the influence of the denominator, which is slightly elevated at onset due to the semi-saturation constant 𝜎^n^. With low contrast, the denominator term is bigger relative to the input drive, resulting in a slower reduced response. For long ISIs, the denominator has sufficient time to decrease in between stimuli, but with shorter ISI, the denominator is still increased due to delayed normalization from the first stimulus, leading to a reduced response to the second stimulus. In both cases, the response rises more slowly and response magnitude is repressed due to a larger denominator term. This figure is produced using *tde_mkFigure6.m*.

At high stimulus contrast, the semi-saturation constant is negligible, and the numerator rises faster than the denominator, leading to a transient (**Fig. 6**, top left). At low contrast, the numerator never rises above the minimum possible value of the denominator, which is set by 𝜎^n^ (**Fig. 6**, bottom left; see Methods and Materials). The main difference between the two contrast conditions thus is the driven response relative to 𝜎^n^. For repeated stimuli, the driven response is effectively the same irrespective of ISI, and the denominator rises and decays similarly for both the first and second stimulus. However, when ISI is long (**Fig. 6**, top right), the denominator has decayed more than when it is short (**Fig. 6**, bottom right). Hence for short ISIs, the ratio of numerator to denominator for the second stimulus is lower. The main difference between the two repetition conditions is thus the level of pre-existing normalization.

In summary, neural responses rise more slowly and are suppressed both when contrast is reduced and when stimuli are repeated with short intervals. According to the DN model, for contrast, this is due to a larger input drive (numerator) than background neural activity (𝜎^n^), while for repetition, the reduction is due to an pre-existing delayed normalization (L\**h_2_*). Thus, the DN model is able to simultaneously predict the effects of both of these stimulus manipulations because it can achieve response reductions through either one of these terms.

### Delayed normalization predicts temporal phenomena throughout visual hierarchy

Next, we examined to what extent each of the canonical computations in the DN model contribute to its ability to predict neural responses for variations in duration, repetition and contrast. To this end, we computed cross-validated explained variance for a number of reduced versions of the model, starting with a linear model with fully parameterized IRFs (see Methods and Materials), after which each of the non-linear operations in the DN model (rectification, exponentiation, normalization, normalization with delay; see **Fig. 1*D***) are added in turn (**Fig. 7*A***). We find that delayed normalization is a key component in raising the prediction accuracy. Adding non-linear operations to a purely linear model gradually increases the ability to explain responses in V1, and a clear gain in explanatory power is observed when adding the final step of delayed rather than instantaneous normalization.

**Figure 7:**
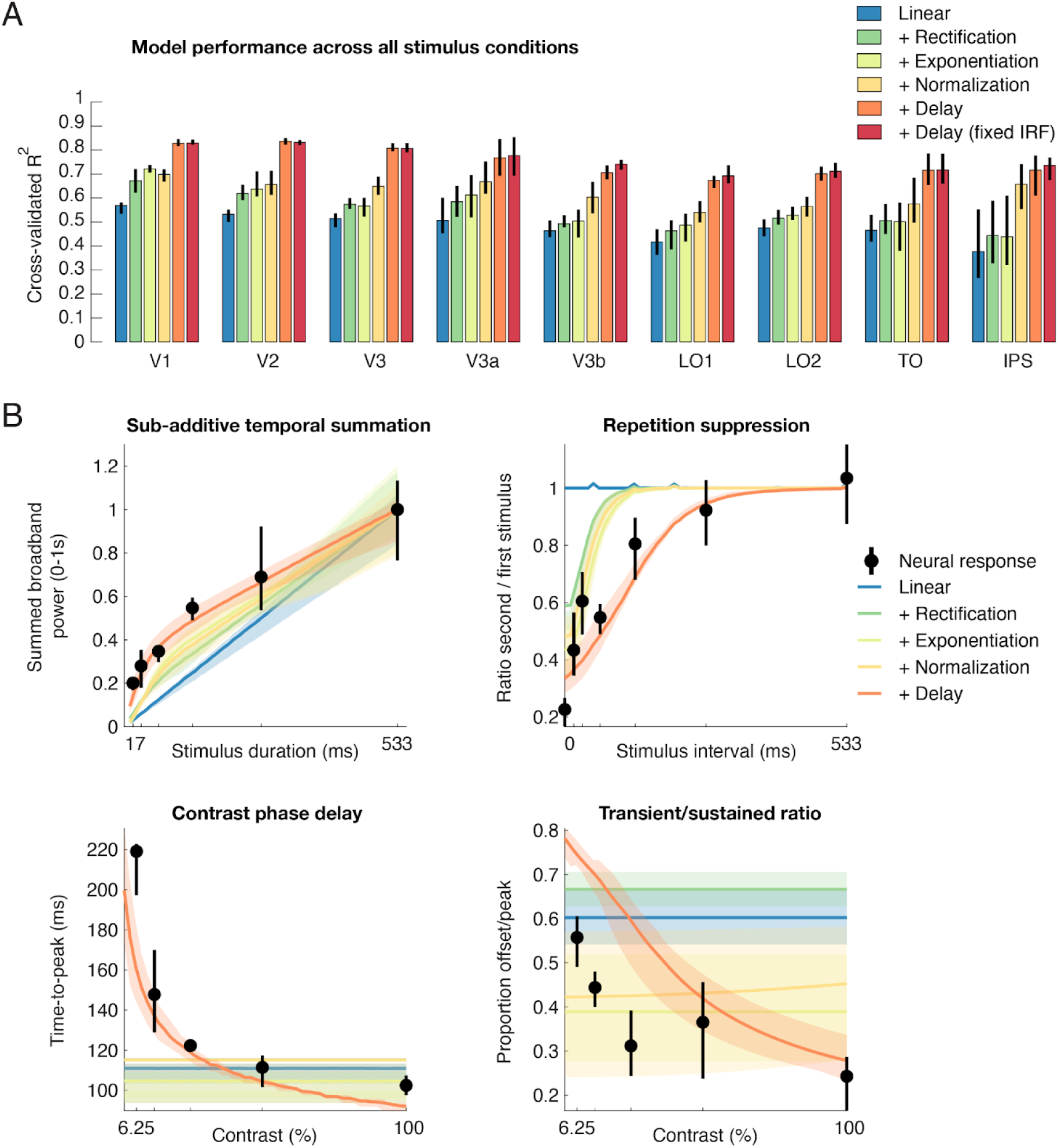
Delayed normalization explains multiple temporal dynamics in multiple visual areas. **A)** Average cross-validated explained variance (coefficient of determination) across all stimulus conditions in V1-IPS for the DN model along with deconstructed versions of the model from which each of the canonical non-linear computations depicted in **Fig. 1*D*** was removed in turn. Relative to a purely linear model (blue), adding non-linear canonical operations of rectification, exponentiation, and delayed normalization each increase the model’s ability to capture the variance in the dataset. Moreover, a full delayed normalized model with a constrained IRF (red) performs equally well as the more unconstrained IRF (2 additional free parameters, orange). Cross-validated explained variance separated by stimulus condition is provided in **Extended Data, Figure 7-1**. **B)** Three non-linear temporal phenomena (sub-additive temporal summation, repetition suppression, and contrast-related dynamics) in V1 as predicted by the deconstructed models. Delayed normalization is especially necessary to capture slowed response dynamics with low stimulus contrast (bottom row). Predicted time courses for each reduced model are provided in **Extended Data Figure 7-2.** Cross-validated model performance and their corresponding temporal phenomena predictions for a set of alternative temporal models (Heeger, 1992; Stigliani et al., 2017, 2019) are provided in **Extended Data, Figure 7-3**. Error bars indicate 68% confidence intervals across 10000 bootstraps of electrode assignments. This figure is produced by *tde_mkFigure7.m*.

Importantly, this pattern of results holds not only in V1, but in all retinotopic maps in which we had sufficient electrode coverage, including V2, V3, and higher-order lateral-occipital and parietal-occipital maps. Because accuracy is computed on left-out data (cross-validation), the result is not guaranteed simply due to adding more free parameters for each model. Indeed, there are some instances where adding parameters makes the fits less accurate (V1, +Exponentiation vs. +Normalization).

In addition, **Fig. 7*A*** shows that the version of the delayed normalization model with fully parameterized IRFs is on par with the more constrained DN model (in which the phase delay and time constants of the negative gamma function are fixed) as used in our analyses so far (see Materials and Methods), indicating that the data do not sufficiently constrain the detailed shape of the IRF, and that the chosen IRF constraints are adequate for the current dataset.

The reduced explanatory power of the deconstructed models is coupled with a reduced ability to predict the different non-linear temporal phenomena induced by our stimulus manipulations (temporal summation, repetition suppression, slower contrast dynamics; **Figs. 2-4**). While compressive temporal summation and repetition suppression can to some extent be captured by reduced models that consist of canonical computations but lack the delayed normalization component (**Fig. 7*B***, top row), delayed normalization is critical to predict the slower dynamics associated with changes in stimulus contrast (**Fig. 7*B***, bottom row). In general, across visual regions, improvements in explained variance were most pronounced for the duration and contrast conditions, with reduced models performing relatively well when predicting responses to stimuli varying in repetition interval (see **Extended Data Figure 7-1** for cross-validated explained variance separated by condition, and **Extended Data Figure 7-2** for time course predictions for each deconstructed model across all stimulus conditions).

### Characterizing temporal dynamics throughout visual hierarchy using the DN model

Having established that the DN model predicts ECoG responses with high accuracy not only in V1, but in multiple visual regions, we next used it to investigate how temporal dynamics change along the visual hierarchy. Note that we use the term “visual hierarchy” as an approximation. By nearly any metric, V2 is a later area than V1, and V3 later than V2, and all the other areas later than V3. However the hierarchical relationship among the areas beyond V3, if any, is uncertain.

### Temporal summation windows increase in higher visual areas

The responses to stimuli that vary in duration revealed a systematic change in temporal summation between visual areas (**Fig. 8**). Compared to V1, responses in, for example, V3b rise more slowly and stay elevated for a longer period of time, resulting in wider responses and higher sustained levels of response relative to the transient (**Fig. 8*A***). We quantified these differences in response shapes across visual areas by computing three different summary metrics, which each capture a different aspect of the response dynamics, on both the data and DN model predictions (see Methods and Materials).

**Figure 8:**
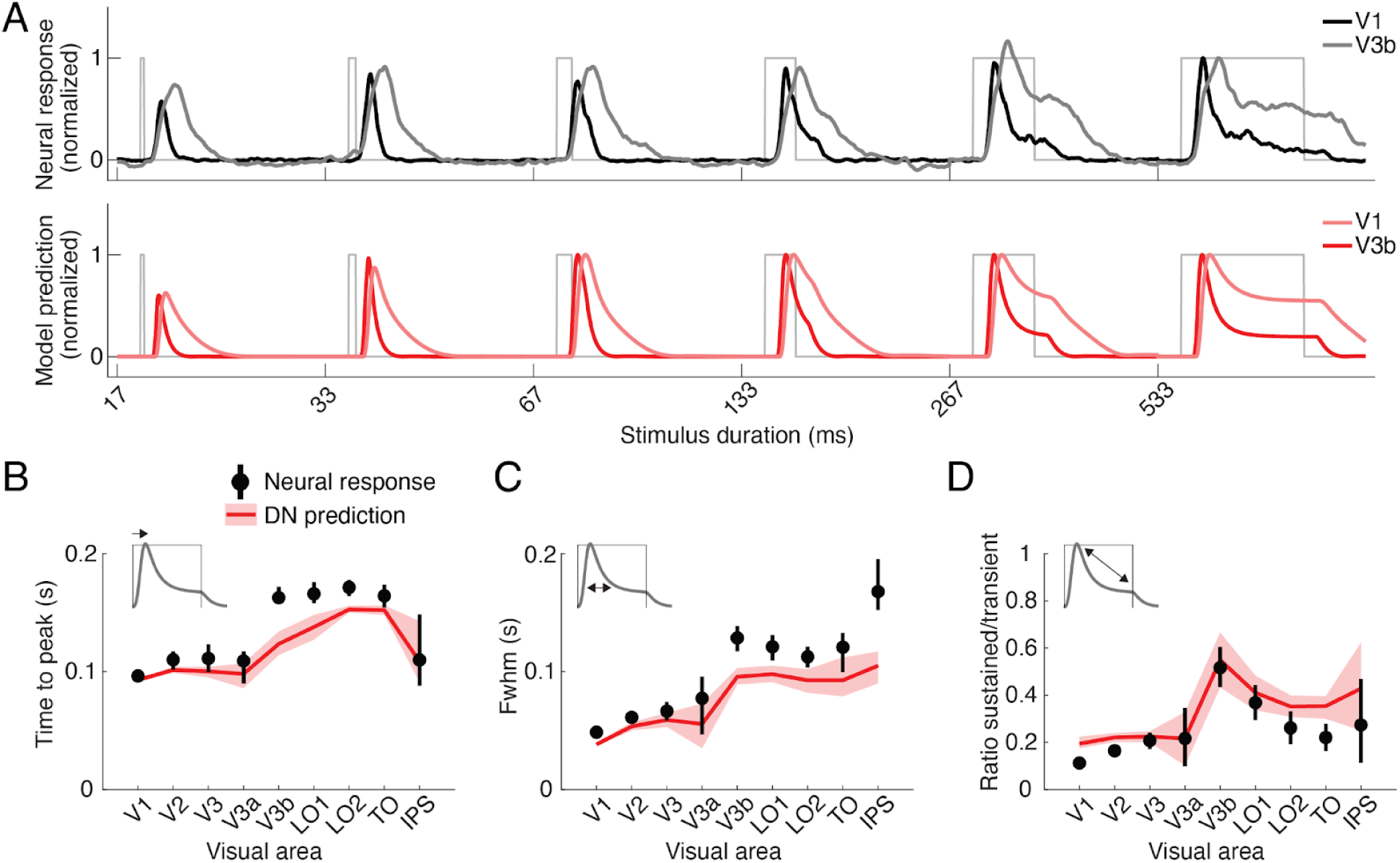
Temporal summation window sizes increase in higher visual areas. **A)** Top: Average broadband ECoG responses for stimuli of increasing duration for an early (V1) and higher visual area (V3b); see **Table 2** for number of included electrodes per area. Bottom: DN model predictions for these same data. Compared to V1, responses in V3b rise more slowly and stay up longer, resulting in wider responses and higher sustained levels of response relative to the transient level. **B-D)** Three summary metrics derived from both the neural responses (black) and DN model predictions (red), separately for all visual areas in the dataset. **B)** Time to peak, computed from the neural response and model prediction to the longest duration stimulus (533 ms). **C)** Full-width half max, computed from the neural response and model prediction to the shortest duration stimulus (17 ms). **D)** Ratio of sustained (response level at stimulus offset) relative to the maximum response (peak), computed from the neural response and model prediction to the longest duration stimulus. Data points indicate means and error bars indicate 68% confidence intervals across 1000 response averages and corresponding DN model fits using probabilistically assigned electrodes. Summary statistics computed separately for individual patients are provided in **Extended Data Figure 8-1**. This figure can be reproduced using *tde_mkFigure8.m*.

Relative to V1, all three measures increased in higher visual areas: the slower rise was reflected in increasing time-to-peak (**Fig. 8*B***); the broadening of the transient was reflected in increasing full width at half max (**Fig. 8*C***); and relatively higher sustained activity was reflected in a sustained/transient ratio that tended to become larger (**Fig. 8*D***), although this latter summary metric was more variable across electrodes compared to the first two metrics. Overall, these summary metrics reflecting differences in temporal summation differ most between a group of earlier visual areas (V1-V3, V3a) and the later areas. These patterns were largely consistent when computed separately within individual participants (see **Extended Data Figure 8-1**).

### Temporal adaptation windows do not differ systematically between visual areas

In contrast to the pattern with temporal summation, adaptation to repeated stimuli showed less evidence of varying systematically across areas (**Fig. 9**). While the response to the second stimulus in V3b is on average higher than in V1 (**Fig. 9*A***), this appears to result from the continued response to the first stimulus even after the second stimulus onset in V3b, rather than to less adaptation (as observed in **Fig. 8**) or a systematically different rate of recovery. In both areas, full recovery was achieved by the longest ISI measured (533 ms). Consistent with these observations, we did not observe a clear relation between the rate of recovery from adaptation across visual areas. To quantify this, we expressed the magnitude of the response to the second stimulus relative to the first stimulus separately for each ISI and visual area, yielding a measure of recovery from adaptation over time per area. While areas V2, V3 and V3a recovered somewhat more slowly than V1 in both the data (**Fig. 9*B***) and DN model predictions (**Fig. 9*C***), area V3b did not. Indeed, when comparing the estimated ISI at which the response is nearly fully recovered (80%) for all areas shows that, for both the data and the model, no systematic change in this measure is observed, with several higher-order visual regions having a similar recovery level as V1 (**Fig. 9*D***).

**Figure 9:**
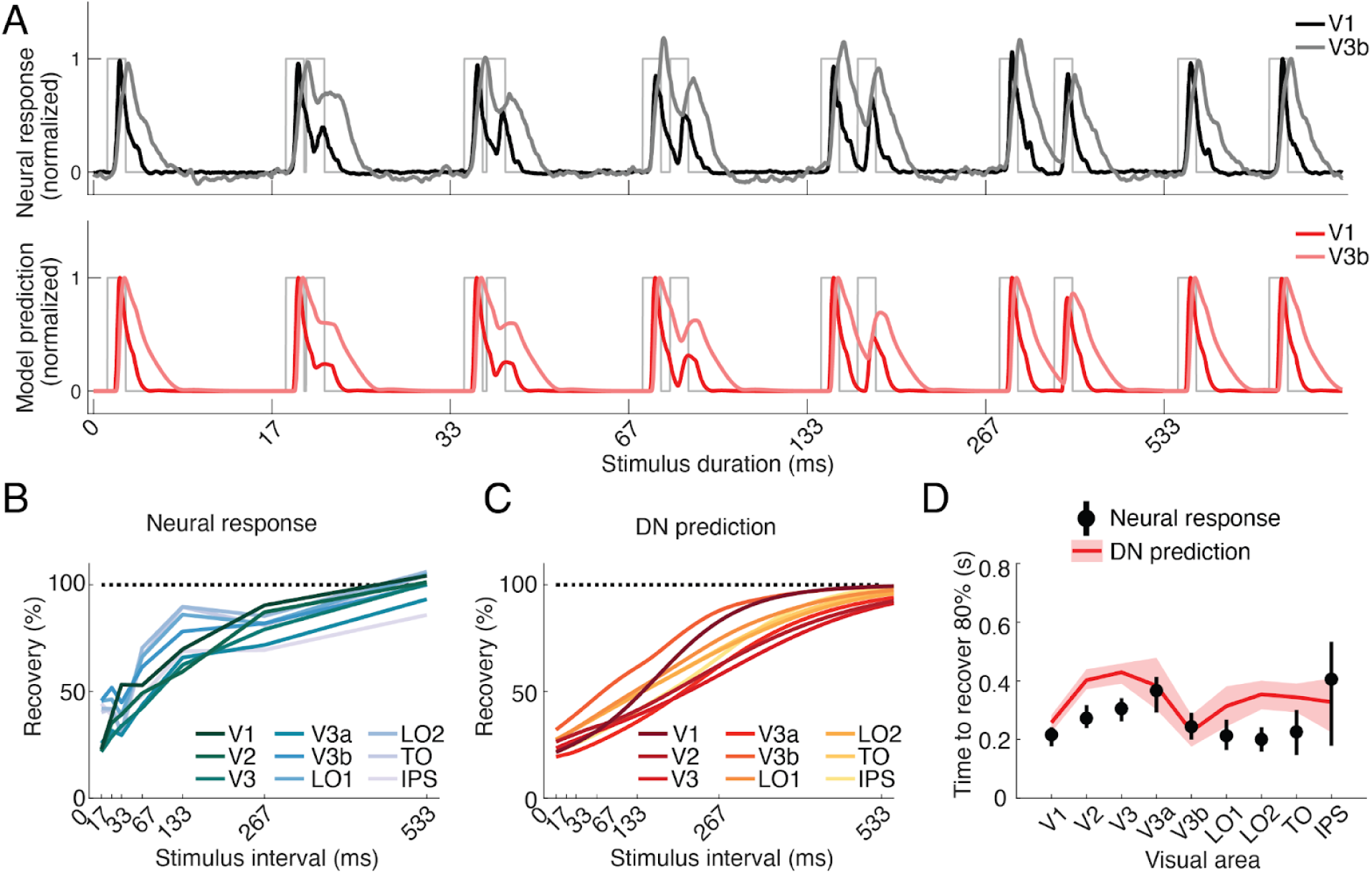
Temporal adaptation windows do not differ systematically between visual areas. **A)** Top: Average broadband ECoG responses to repeated stimuli with increasing ISI for an early (V1) and a higher visual area (V3b); see **Table 2** for number of included electrodes per area. Bottom: DN model predictions corresponding to these same data. Compared to V1, responses in V3b recover at a similar rate, with full recovery being achieved at the longest ISI measured (533ms). **B-D)** Recovery from adaptation across visual areas. While most higher visual areas show a somewhat slower recovery compared to V1 (e.g. V3a) some areas show a similar or faster recovery (e.g. V3b, LO1) and error bars are highly overlapping, suggesting no systematic change in recovery across the visual hierarchy. **B)** Measured recovery from adaptation across ISIs for V1-V3b (calculated the same way as in **Fig. 3*B***). **C)** Predicted recovery from adaptation based on the DN model when simulating responses across a range of ISIs. **D)** Estimation of the ISI at which recovery is 80% of the response to the first stimulus, calculated for both neural data and DN model. Data points indicate means and error bars indicate 68% confidence intervals across 1000 response averages and corresponding DN model fits using probabilistically assigned electrodes. This figure is produced using *tde_mkFigure9.m*.

### Opposite effects of contrast on response amplitude and latency across visual areas

In a third comparison of visual areas, we examined responses as a function of contrast (**Fig. 10**). Interestingly, we observed opposite effects of contrast on response amplitude and timing across visual areas. Amplitude differences between low and high contrast stimuli become smaller in higher-visual areas (greater invariance to contrast), whereas time-to-peak differences tend to get larger (greater sensitivity to contrast).

**Figure 10:**
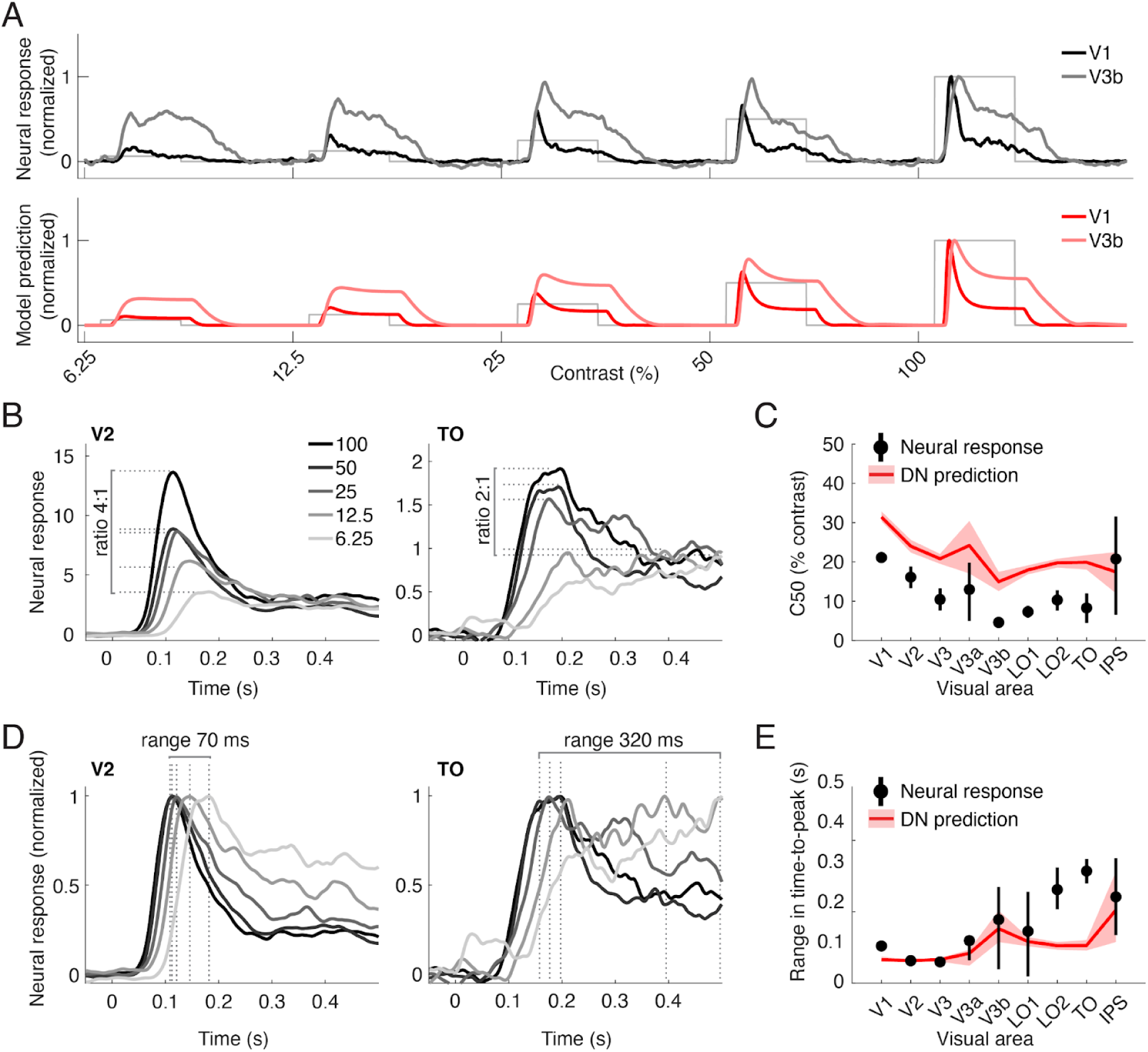
Opposite effects of contrast on response amplitude and timing across visual areas. **(A)** Top: Average broadband ECoG responses to stimuli with increasing contrast for an early (V1) and a higher visual area (V3b); see **Table 2** for number of included electrodes per area. Bottom: DN model predictions corresponding to these same data. **B)** Average broadband ECoG responses to stimuli with different contrasts in early visual area V2 (left) and higher visual area TO (right), superimposed. Lines indicate maximum amplitude in each condition. Time courses were slightly smoothed for illustrative purposes only. **C)** C50 for each visual area, computed both from the data and model predictions. **D)** Same data as in B), with each time course normalized to its peak, which illustrates differences in response rise. Lines indicate time-to-peak in each condition. **E)** Range in time-to-peak across contrast for each visual area for both data and DN model predictions. Data points in C) and E) indicate means and error bars indicate 68% confidence intervals for 1000 response averages of probabilistically assigned electrodes. This figure is produced using *tde_mkFigure10.m*.

The amplitude effects can be seen by comparing responses across different contrast conditions within a given area: compared to V1, responses in V3b are relatively enhanced for low contrast stimuli (**Fig. 10*A***); similarly, compared to V2, responses in TO are relatively enhanced for low contrast stimuli (**Fig. 10*B***). We quantified this pattern by fitting a Naka-Rushton equation to the peak responses for both the data and the DN model predictions (see Methods and Materials). The C50 parameter is the contrast at which the response reaches half its maximum, with lower values indicating steeper rises in contrast response functions. This measure showed a downwards trend along the visual hierarchy in both the data and the model predictions (**Fig. 10*C***), indicating that less contrast is needed to elicit a robust response in higher compared to lower visual areas. While the data and model predictions agree in showing decreasing C50 in higher areas, there is an overall offset between data and model, with the model predicting slightly higher C50 parameters overall. This matches our previous observation of small but systematic underpredictions of the transient response at low contrast noted in **Fig. 4*C***.

Differences in the time to peak between low and high contrast stimuli are more pronounced in the highest visual areas. This is illustrated in **Fig. 10D**, which shows the same data as in **Fig. 10B** but with each condition normalized to its peak (see **Fig. 4A**, right panel for same data in V1). Both V2 and TO show a greater and shallower slope at lower contrast (**Fig. 10D**). However, in area TO the range in latencies is larger compared to V2. To quantify these effects, we calculated the time-to-peak for each contrast (i.e., the dashed vertical lines in **Fig. 10D**) and measured the range (difference between minimum and maximum value) across contrast levels in each area, for both the data and the model (**Fig. 10E**). While the range is relatively constant in early areas V1-V3, both data and model show an upwards trend in the time-to-peak range across contrast conditions in the highest visual areas. This suggests that, unlike response amplitude, response latency becomes more sensitive to contrast in higher visual areas.

### Changes in temporal dynamics across visual areas

To summarize the comparisons across areas, we find that the temporal dynamics of neural responses differ across the visual areas we measured in several different ways. Most changes seem to reflect increasing invariance in higher visual areas, demonstrating, for example, less dramatic effects of changing stimulus duration on neural response shapes, or less effect of changing contrast intensity on response amplitude, compared to lower visual areas. However, not all of our analyses were indicative of increased invariance along the visual hierarchy. In particular, recovery from adaptation remained fairly stable, suggesting that the rate of adaptation does not change systematically across areas, or that our measurements were not sensitive enough to pick up differences in this response property. (We return to this in the Discussion). In addition, response latency becomes more sensitive to stimulus contrast in higher visual areas. Importantly, changes in response properties between low and higher visual areas were to a large extent recapitulated by the DN model, suggesting that the model captures temporal response dynamics not just in early visual regions but across visual cortex more broadly.

## Discussion

We demonstrated several non-linearities in the neural dynamics of visual cortex: saturation as stimulus contrast and duration increase, suppression for repeated stimuli at short intervals, and slower onsets at low contrast. While each of these has been demonstrated previously (e.g., Albrecht and Hamilton, 1982; Albrecht et al., 2002; Zhou et al., 2018), an important goal in systems neuroscience is to understand how apparently disparate phenomena might be linked (Wandell et al., 2015). Here we showed that a delayed normalization model predicts these responses in multiple human retinotopic maps.

### Fitting one model to many stimuli

This DN model was developed to account for similar temporal phenomena in separate, independent studies (Zhou et al., 2019). While that work showed that a single model *form* could account for a wide variety of neural data, the various datasets were fit separately because they came from different experiments, measurement types, and species. Temporal dynamics and fitted parameters differed substantially across datasets: for example, the time to peak of single unit macaque V1 data was tens of ms, but over 100 ms for human ECoG broadband. A key advance here is that, for a given cortical site, a single instance of the DN model fit *simultaneously* to many stimulus types captures temporal dynamics arising from temporal summation, adaptation, and variation in contrast.

### Parallels between responses to repeated and low contrast stimuli

A surprising observation is that response time courses to a repeated stimulus at high contrast and a non-repeated stimulus at low contrast are remarkably similar. The delayed normalization model accounts for this through the time-varying relationship between stimulus drive and normalization: At low contrast, the stimulus drive never gets large relative to the normalization, and for repeated stimuli, there is lingering normalization from the first stimulus.

The similarity in responses does not imply that the two stimulus dimensions are perfectly conflated in the brain. For example, monkeys can detect stimulus repetitions irrespective of contrast, implying some contrast-independent stimulus memory (Mehrpour et al., 2020). Nonetheless, the parallels in ECoG responses raise the prospect that behavioral measures, for example detection thresholds or response time, will show similar patterns for the two stimulus manipulations, and might be explained by a single model. Such a model could clarify how stimulus contrast interacts with cognitive phenomena like the attentional blink (Raymond et al., 1992). A dynamic normalization model has recently been used to explain behavioral effects of temporal attention (Denison et al., 2021). Our findings suggest that a dynamic normalization model might also explain how effects of temporal attention interact with stimulus contrast.

### Links between ECoG, fMRI and single-unit electrophysiology

We quantified ECoG responses as the time-varying broadband envelope. Field potential results differ dramatically depending on the measure. For example, the broadband response, but not the steady state evoked potential, shows subadditive spatial summation (Winawer et al., 2013); Narrowband gamma oscillations and broadband power elevations show different stimulus selectivity (Ray and Maunsell, 2011; Hermes et al., 2015, 2019). We studied the broadband response because it appears most strongly connected to other measures, including fMRI (Hermes et al., 2017), spiking (Miller et al., 2009; Ray and Maunsell, 2011), and behavior (Miller et al., 2014), thereby facilitating comparisons with those literatures.

The temporal phenomena we report have been demonstrated with fMRI and single-unit physiology. However, these methods leave open some questions better addressed with ECoG. For example, fMRI visual responses show sub-additive temporal summation (Boynton et al., 1996; Huettel and McCarthy, 2000; Birn et al., 2001; Mukamel, 2004; Zhou et al., 2018). These effects could reflect a mixture of non-linear summation in the BOLD signal and neural activity. Here, we demonstrated temporal subadditivity in population responses in human ECoG, paralleling single-cell recordings (Tolhurst et al., 1980). Similarly, fMRI shows response suppression to repeated stimuli (Grill-Spector et al., 2006). However, responses to stimuli separated by short intervals cannot be measured independently with fMRI, whereas ECoG allows for separate estimations of the response time courses. Mirroring single-cell recordings (Motter, 2006), we find that adaptation is associated with both a reduction of response amplitude and slower rise time.

The ECoG measures also provide important data not easily obtained in animal studies. We made systematic comparisons of temporal dynamics across many visual areas, which is typically not feasible in microelectrode recordings.

### Changes in temporal dynamics across visual areas

Several studies find that time-scales of temporal processing become longer ascending the visual hierarchy (Hasson et al., 2008; Weiner et al., 2010; Honey et al., 2012; Mattar et al., 2016; Zhou et al., 2018), paralleling increases in spatial receptive fields (Maunsell and Newsome, 1987). In contrast, a recent fMRI study reported no differences in the recovery time of repetition suppression across visual areas (Fritsche et al., 2020). We replicated both patterns, suggesting there is not a single processing time-scale per area. When temporal dynamics are characterized by the period over which responses sum, we find systematic increases in time scale from earlier to higher areas (Figure 8). However, when characterized as recovery time from adaptation (Figure 9), time scales were relatively constant, consistent with Fritsche et al., (2020). These patterns are not contradictory, as both are also observed in the model fits, indicating that even a relatively simple model can result in complex behavior.

Another property that varies across visual areas is contrast sensitivity (e.g,. Tootell et al., 1998; Avidan et al., 2002). As with temporal windows, the degree of contrast invariance depends on the quantification. Specifically, response amplitudes become more invariant in higher visual areas, but response latencies do not; if anything, higher areas show greater latency differences. Thus, some information about stimulus contrast remains in the neural responses of later areas. The fact that sensitivity to contrast remains in the response timing but not amplitude means that measurements which average across a trial (like fMRI or mean spike rates) will miss this feature.

### Space and time

Some changes in dynamics across visual areas parallel findings in the spatial domain, for example spatial receptive fields and temporal summation windows increase in higher areas. However, spatial and temporal properties are not perfectly separable. To make analyses tractable, we kept spatial stimulus properties (other than contrast) constant. Our stimuli effectively drive responses in multiple visual maps, especially V1-hV4 (Kay et al., 2013b; Zhou et al., 2018), but higher maps may respond better to more complex stimuli (Sayres and Grill-Spector, 2008; Arcaro et al., 2009; Silson et al., 2016). We recently found that ventral-temporal and lateral-occipital ECoG responses were several times larger for naturalistic images than for simple textures, with the amount of adaptation depending on the electrode’s category preference (faces, bodies, objects; Brands et al., 2021). A complete characterization of temporal dynamics thus also requires incorporating spatial stimulus properties (and *vice versa*). An important yet unachieved goal for the field is a space-time model that simultaneously accounts for spatial summation, stimulus selectivity and temporal dynamics throughout visual cortex.

### Model failures and limitations

While the DN model captured many response variations, we also noted some small but systematic model failures. The model underpredicts the transient response at low contrasts and for short ISI repeats. The DN model produces response transients because of sluggish normalization, whereas other models account for transient and sustained responses with distinct temporal channels (Horiguchi et al., 2009; Stigliani et al., 2017). It is likely that neither type of model is complete. While multiple-channel models do not explain slower dynamics at low contrast (see **Extended Data Figure 7-3**), incorporating multiple channels into the driven responses of a DN model might help ameliorate model failures at low contrast and short ISIs.

Our results are subject to several limitations. First, not all participants contributed data to all maps. When differences in response properties are expected to be small (say, V1 vs. V2), variability across participants may be larger than the map differences. Therefore, our comparisons focused primarily on coarse groupings: earlier (V1-V3) versus later maps. Second, we did not measure retinotopic maps in individual participants leading to some uncertainty in electrode assignments. For this reason, we developed a probabilistic resampling method to incorporate this uncertainty into our measures of variability. Third, we did not have electrodes with sufficiently reliable responses in ventral stream regions beyond V3. Those regions could exhibit different temporal dynamics than lateral-occipital and dorsal-parietal maps (Stigliani et al., 2019), that may or may not be well captured by the DN model. An important future direction is to measure temporal dynamics in neural population responses with more extensive sampling of ventral visual cortex.

The fact that the same model form accurately fits responses in several cortical areas does not imply that each area is computing the same thing. Rather, it implies that some aspects of the responses are captured by a common model form (though differing in parameters). There is in fact evidence that cortical areas with very different stimulus selectivity, V1 and MT, manifest a similar computational architecture, differing primarily in their inputs rather than computations (Heeger et al., 1996). Fitting a single model form (such as a spatial receptive field) with separate parameters to different locations in the visual pathways facilitates comparisons. More generally, our model of any area, say V1, is not a model of what that area computes, but rather a model that summarizes the computations of the complete circuitry from eye to brain, including feedforward, lateral, and feedback pathways, sampled at that location. Some of the normalization measured in the V1 responses may be inherited from earlier processing stages, or arise from feedback from higher areas. This is the same logic used when, say, measuring the receptive field of a visual neuron in terms of visual field locations.

Finally, this dataset may serve as a useful benchmark for testing other models. As an example, we computed predictions for our data from a few previously published models (Heeger, 1992; Stigliani et al., 2017, 2019) (**Extended Data Figure 7-3**). To facilitate model comparison, we make the data publicly available with open code that implements models modularly, allowing fitting multiple models to the same data using automated cross-validation and area comparison.

## Acknowledgements

This work was funded by BRAIN Initiative, the National Institute Of Mental Health of the National Institutes of Health under Award Number R01MH111417. The content is solely the responsibility of the authors and does not necessarily represent the official views of the National Institutes of Health.

## Extended Data

**Extended Data Figure 1-1:**
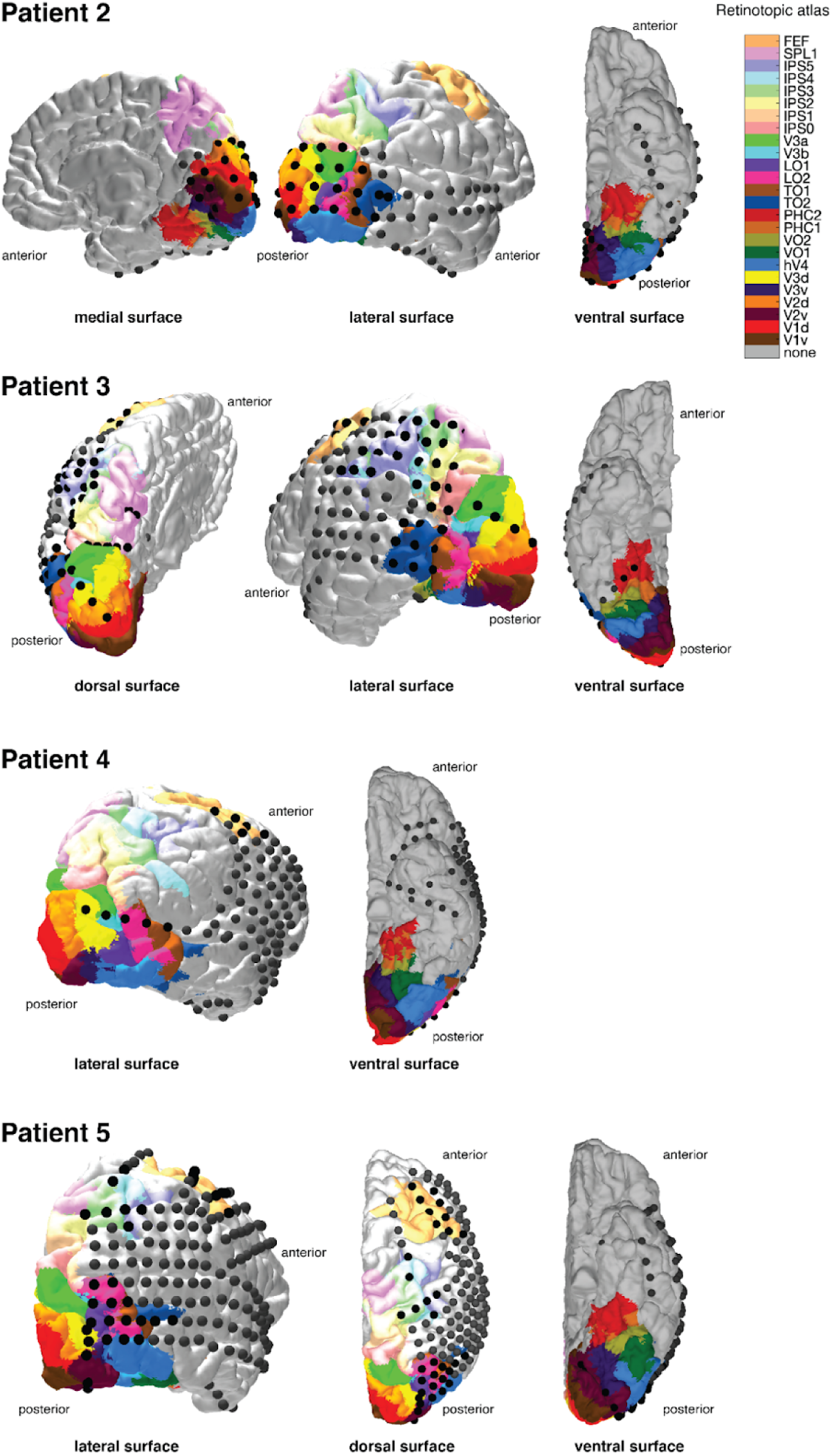
Electrode reconstruction on pial surfaces of patient 2, 3, 4 and 5. Different views of the implanted hemisphere(s) are shown and labeled for each patient. The normalized full probability atlas (see Materials and Methods) (from Wang et al., (2014) is projected on each patient’s pial surface and depicted using the color legend shown in the upper right. 3D interactive images of the individual meshes can be reproduced using *tde_mkFigure1_1.m*

**Extended Data Figure 1-2:**
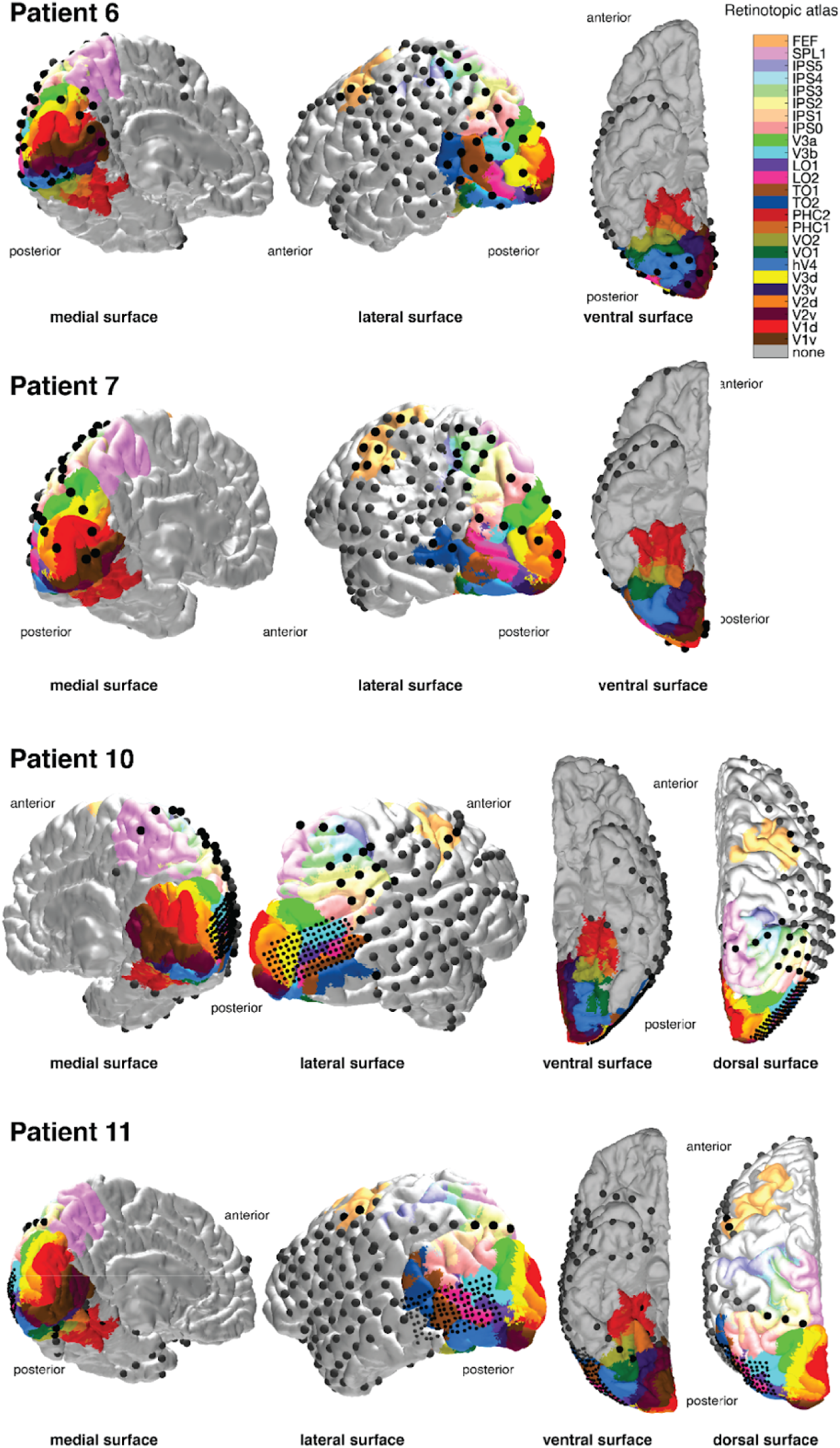
Electrode reconstruction on pial surfaces of patient 6, 7, 11 and 11. Same as Extended Data Figure 1-1, but for patients 5-8. 3D interactive images of the individual meshes can be reproduced using *tde_mkFigure1_1.m*

**Extended Data Figure 1-3:**
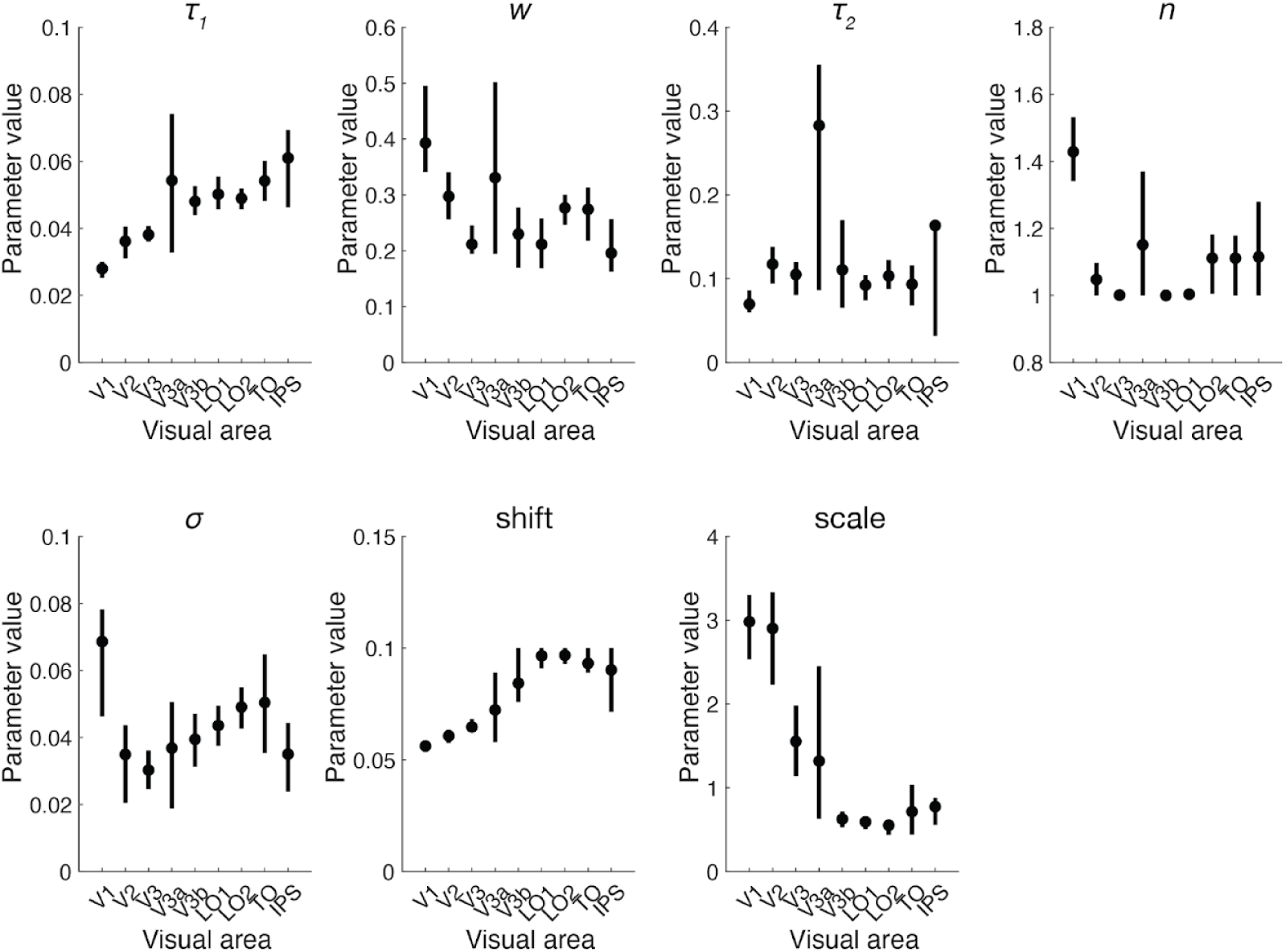
DN model parameters for all visual areas. The DN model has five model parameters that were fitted across all stimulus conditions: 𝜏_1_ (time constant of the IRF), w (weight of the negative and positive IRFs), n (exponent), 𝜎 (semi-saturation constant), and 𝜏_2_ (time constant of the exponential decay), as well as two nuisance parameters: shift (delay in response onset relative to stimulus onset) and scale (gain of the response). Data points indicate median across probabilistically assigned electrodes. Error bars indicate 68% confidence intervals across 10,000 bootstraps of electrode assignments to visual areas. Initialization parameter values and bounds used during fitting can be found in json meta-data files accompanying the model code provided in the folder ‘temporal_models’ in https://github.com/irisgroen/temporalECoG. This figure can be reproduced using *tde_mkFigure1_3.m*.

**Extended Data Figure 7-1:**
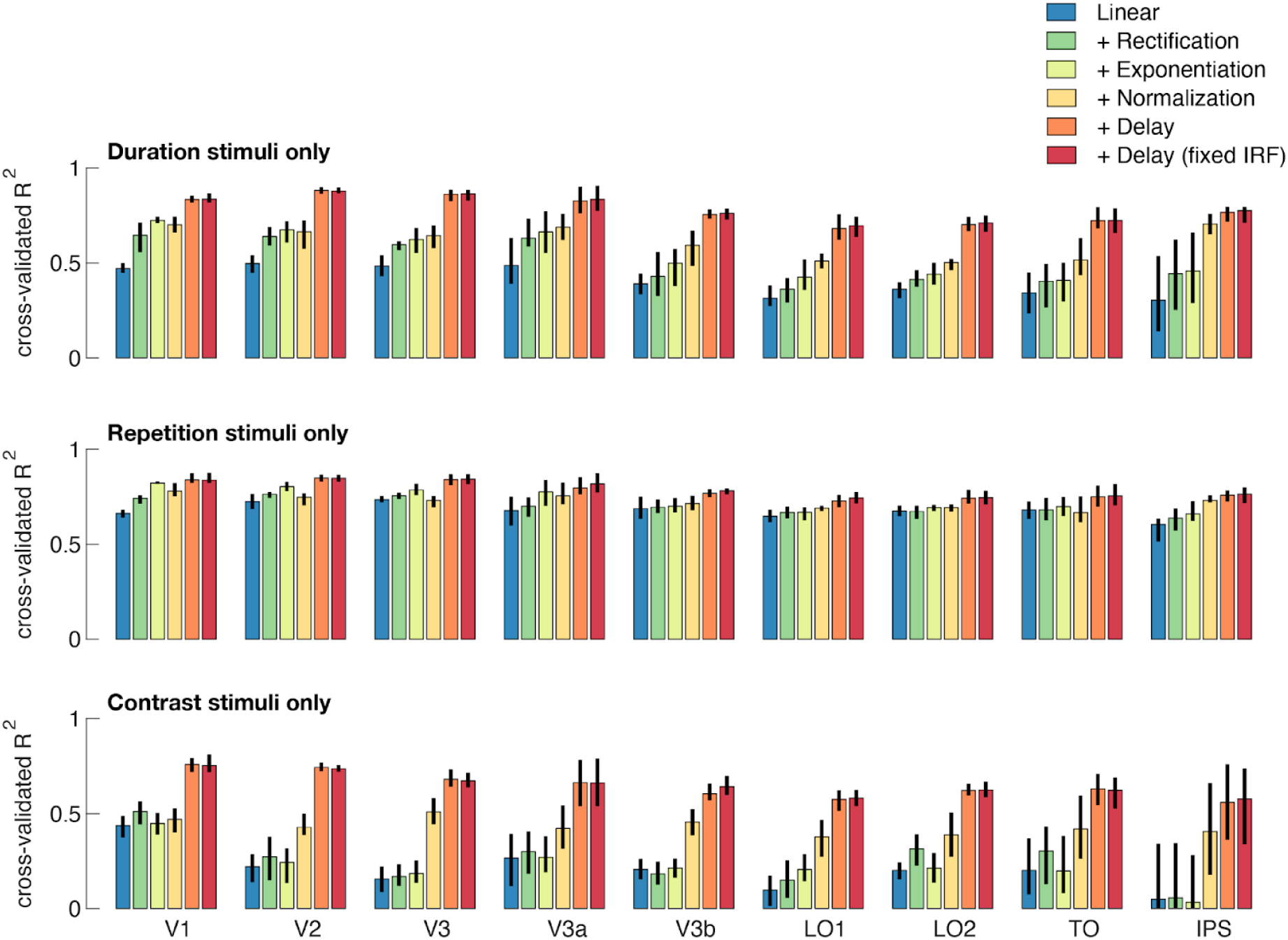
Variance explained by DN model, separated by stimulus condition. Cross-validated R2 DN model components, separated by stimulus manipulation. Note that the models were fit to all conditions simultaneously, but that model fits here are evaluated separately for the duration, repetition and contrast conditions. Error bars indicate 68% confidence intervals across 10000 bootstraps of electrode assignments. This figure is produced by *tde_mkFigure7_1.m*.

**Extended Data Figure 7-2:**
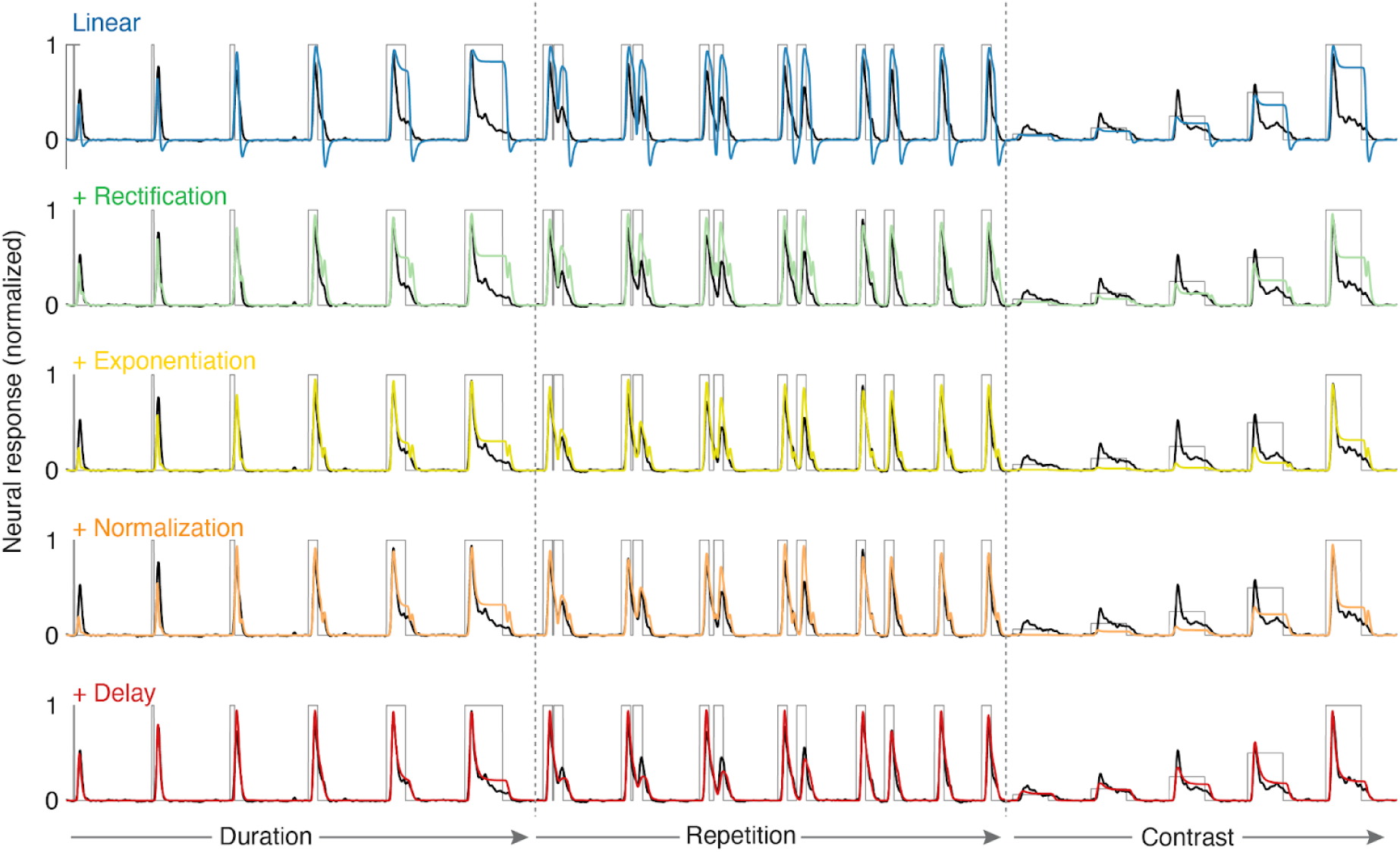
Predicted time courses for deconstructed DN model. V1 time-courses with deconstructed DN model predictions from Figure 7 overlaid, for all stimulus types (duration, repetition, contrast). A linear model consisting of only a biphasic IRF clearly fails to explain the neural responses across conditions. Adding more canonical computations, including instantaneous normalization, results in increased predictive performance in capturing the overall neural response shape, in particular for the duration and repetition conditions. Adding a temporal delay in normalization is especially critical for capturing changing temporal dynamics (e.g. time-to-peak, sustained/transient ratio) for contrast-varying stimuli. Data and model time courses were averaged across electrodes using a 1000 bootstraps of electrode assignments. This figure can be reproduced using *tde_mkFigure7_2.m*.

**Extended Data Figure 7-3:**
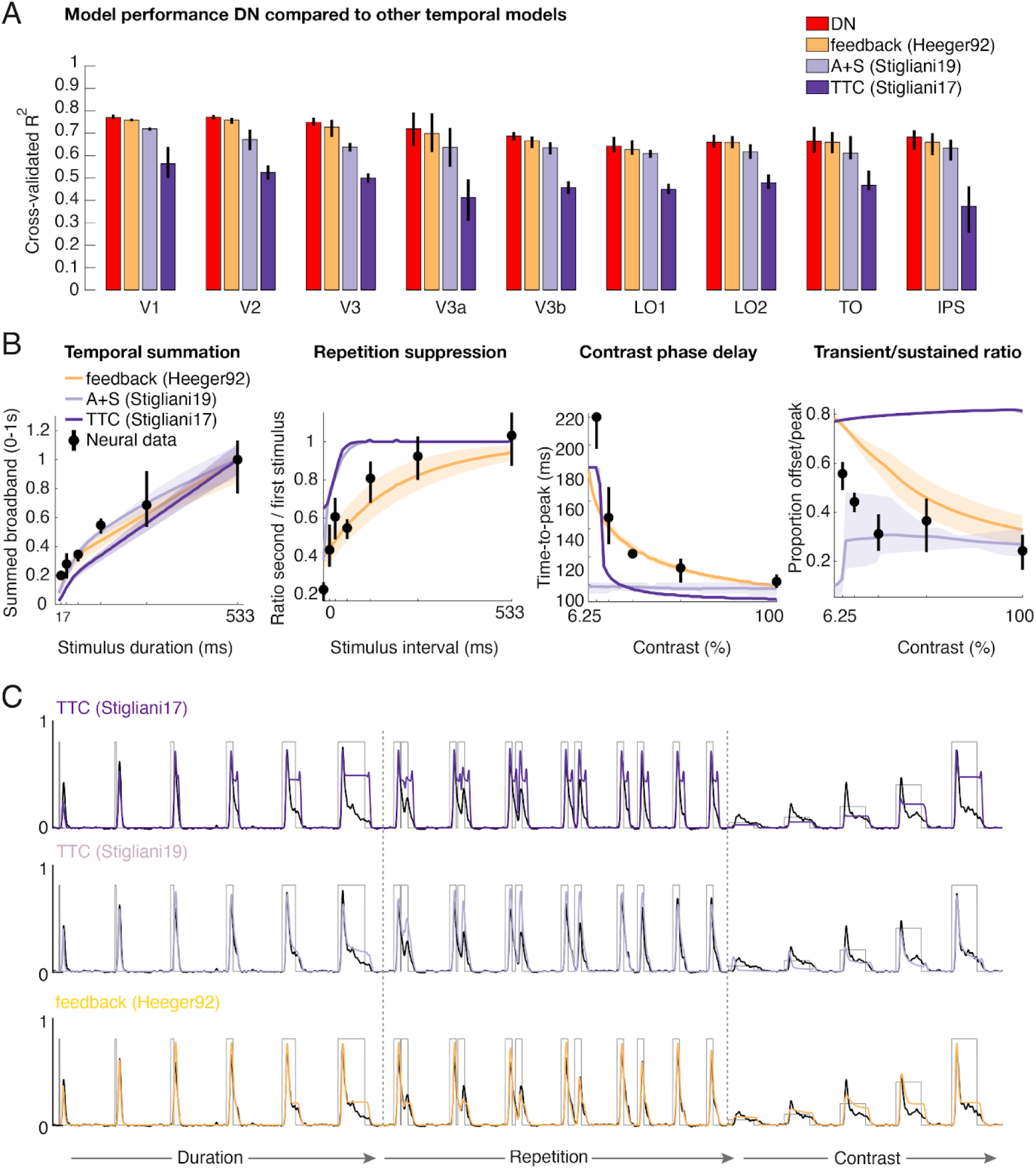
Alternative model performance and predictions. **A)** Cross-validated explained variance in V1-IPS for the DN model compared to a model implementing history-dependent normalization via feedback (Heeger, 1992) and two 2-channel (A+S, TTC) models (Stigliani et al., 2017, 2019). **B)** Non-linear temporal phenomena in V1 (same data as in **Fig. 7*B***) as predicted by the alternative models. **C)** V1 time-courses with alternative model predictions overlaid, for all stimulus types. Two-temporal channel models can predict temporal summation and repetition suppression effects to some degree, but fail to capture contrast-induced temporal dynamics. In contrast, the feedback model performs on par with the DN model. This figure can be reproduced using *tde_mkFigure7_3.m*.

**Extended Data Figure 8-1:**
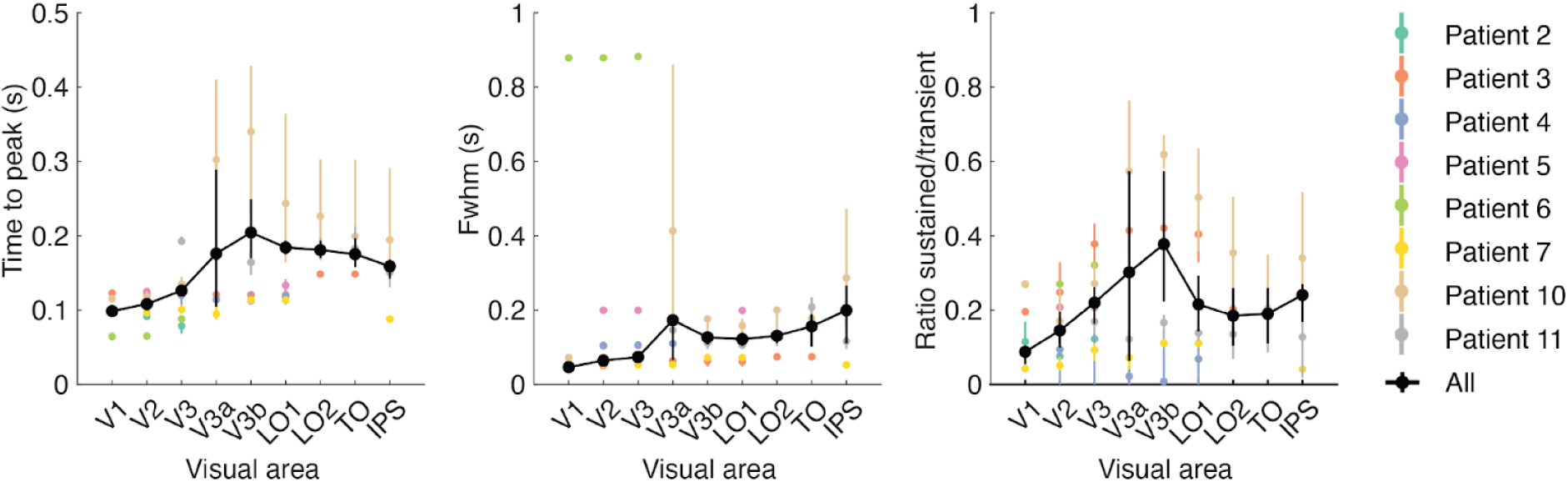
Temporal summation windows for individual participants. Time to peak, Fwhm and Sustained/transient ratio computed from the data, but separately for each electrode and then grouped by visual area within each individual patient (colored data points) as well as across all patients (black data points and line). Note that the average here is not identical to the data shown in **Figure 8*B-D*** where summary statistics were computed based on average time courses per area, while here they were calculated per individual electrode and then averaged. Data points indicate median across repeated samples of electrodes, taking into account each electrode’s probability of overlap with a retinotopic atlas (see Methods). Error bars and shaded regions reflect 68% confidence intervals across 1,000 bootstraps of electrode assignments. While individual electrode data is noisy, and there is substantial variety amongst individual patients, most patients show trends towards longer temporal summation windows for higher visual areas, as evidenced by increase in time-to-peak, response width and sustained to transient ratios. This figure can be reproduced using *tde_mkFigure8_1.m*.

